# Dating ammonia-oxidizing bacteria with abundant eukaryotic fossils

**DOI:** 10.1101/2024.02.23.581699

**Authors:** Tianhua Liao, Sishuo Wang, Hao Zhang, Eva E. Stüeken, Haiwei Luo

## Abstract

Evolution of a complete nitrogen cycle relies on the onset of ammonia oxidation, which aerobically converts ammonia to nitrogen oxides. However, accurate estimation of the antiquity of ammonia-oxidizing bacteria (AOB) remains challenging because AOB-specific fossils are absent and bacterial fossils amenable to calibrate bacterial molecular clocks are rare. Leveraging the ancient endosymbiosis of mitochondria and plastid, as well as using state-of-the-art techniques such as the Bayesian sequential dating approach, we obtained a robust timeline of AOB evolution calibrated by fossil-rich eukaryotic lineages. We show that the first AOB evolved in marine Gammaproteobacteria (Gamma-AOB) and emerged between 2.1 and 1.9 billion years ago (Ga), thus postdating the Great Oxidation Event (GOE; 2.4-2.32 Ga). To reconcile the sedimentary nitrogen isotopic signatures of ammonia oxidation occurring near the GOE, we propose that ammonia oxidation likely occurred at the common ancestor of Gamma-AOB and Gammaproteobacterial methanotrophs, or the actinobacterial/verrucomicrobial methanotrophs, which are known to have ammonia oxidation activities. We also do not rule out another possibility that nitrite was transported from the terrestrial habitats where ammonia oxidation by archaea likely took place. Further, we show that the Gamma-AOB predates the anaerobic ammonia oxidizing (anammox) bacteria which also emerged in marine environments, implying that the origin of ammonia oxidation constrained the origin of anammox as nitrite produced by the former fuels the latter. Our robustly calibrated molecular clocks support a new hypothesis that nitrogen redox cycle involving nitrogen oxides evolved rather late in the ocean.

## Introduction

The nitrogen (N) cycle encompasses a series of transformations between ammonia, dinitrogen, and nitrogen oxides, mediated by various biological processes such as dinitrogen (N_2_) fixation and nitrification (Preena et al. 2021; Pajares and Ramos 2019; Yool et al. 2007; Kuypers et al. 2018; Stüeken et al. 2016). The evolutionary onset of these processes, especially nitrification (i.e., the oxidation of ammonia to nitrite and nitrate), has important implications for biological activity and other biogeochemical cycles. The production of nitrogen oxides enables biological or abiotic denitrification (i.e., reduction of nitrate and nitrite to N_2_), leading to a depletion of the dissolved N reservoir. It has therefore been hypothesized that the onset of nitrification and denitrification resulted in a N-limited biosphere, which may have impacted microbial evolution (Fennel et al. 2005). Reconstructing the emergence of nitrification in the biosphere is therefore critical for understanding the evolutionary history of life on Earth.

Geochemical techniques, specifically stable N isotopes (δ^15^N) applied to organic-rich sedimentary rocks, provide an indirect tool for constraining biological metabolisms. These data suggest that a dissolved reservoir of nitrate appeared transiently in the Neoarchean surface ocean ca. 2.7 billion years ago (Ga), and the most plausible way to generate dissolved nitrate in significant quantities is by biological nitrification (Koehler et al. 2018; Godfrey and Falkowski 2009). These data are therefore used as indirect evidence for local and/or temporary occurrences of oxygenic photosynthesis. The δ^15^N signal becomes more widespread around the time of the Great Oxidation Event (GOE, 2.4 to 2.32 Ga) (Zerkle et al. 2017; Kipp et al. 2018; Cheng et al. 2019) which marks the permanent oxygenation of the atmosphere and surface environment. The interpretation of the δ^15^N record is consistent with other proxies; however, it is by itself not unambiguous. First, post-depositional processes, like diagenesis and metamorphism, can alter sedimentary proxies, which creates uncertainty (Robinson et al. 2012; Ossa et al. 2022; Stüeken et al. 2016). Second, and perhaps more importantly, the δ^15^N signal interpreted as evidence of coupled nitrification/denitrification is non-unique and may be mimicked by other metabolisms (e.g., distillation of a dissolved ammonium reservoir by partial assimilation or Fe-based oxidation; (Pellerin et al. 2023)). Therefore, new techniques are needed to provide independent constraints. Here we leverage the genomic record of the biosphere, paired with bioinformatic reconstructions of the antiquity of nitrifying bacteria, to determine the age of bacterial nitrification.

The nitrification process is mediated by several microbial functional groups, including the ammonia-oxidizing archaea (AOA) and bacteria (AOB), the nitrite-oxidizing bacteria (NOB) (Park et al. 2020; Zorz et al. 2018; Lehtovirta-Morley 2018), and complete ammonia-oxidizing bacteria (CAOB or Comammox bacteria) (van Kessel et al. 2015). The bacterial ammonia oxidizers are limited to a few taxa, including members of Beta- (Beta-AOB) and Gammaproteobacteria (Gamma-AOB) and the genus *Nitrospira* within the phylum Nitrospirota. Although taxonomically distinct, they use the same enzyme for ammonia oxidation, i.e., ammonia monooxygenase (AMO), which is encoded by the gene cluster *amoCAB* and belongs to the copper membrane monooxygenase family (CuMMO). The genes encode the CuMMO monooxygenases, regardless of the substrate specificity, are referred to as *xmoCAB* (Tavormina et al. 2011; Khadka et al. 2018). Genome-scale analysis focusing on auxiliary genes helps to identify the potential substrate preference of *xmoCAB*-containing bacteria. For example, bacterial ammonia oxidizers utilize the ubiquinone redox module (HURM) (Simon and Klotz 2013; Klotz and Stein 2008) for energy harvesting. This module requires the presence of two electron carriers, soluble cytochrome *c554* and cytochrome *c_M_552* encoded by the auxiliary genes *cycAB* (Simon and Klotz 2013). Interestingly, pure culture-based studies reported the ability of *pmoCAB*-containing but *cycAB*-lacking bacteria to oxidize ammonia (Jones and Morita 1983; O’neill and Wilkinson 1977). Because of this substrate promiscuity, the antiquity of ammonia oxidation, if solely estimated based on the *amoCAB* sequences or *amoCAB*-containing microbes, is likely underestimated.

Several studies attempted to date the origin of ammonia oxidizers using molecular clock methods, which assume that the evolutionary rates of protein sequences in different lineages were approximately constant (Zuckerkandl et al. 1962) and employ relaxed molecular clock models to account for rate variations across lineages (Dos Reis et al. 2016; Dos Reis et al. 2018). Despite great efforts, the results of previous studies seem often contradictory. Specifically, Ren et al. (2019) estimated that AOA emerged in terrestrial habitats at ∼2.1 Ga and they expanded to oceanic niches at ∼1.0 Ga. A later study (Yang et al. 2021), while agreeing on the ocean-terrestrial habitat transition, suggested a much younger origin of AOA at 1.2 Ga. Notably, both studies used secondary calibrations (i.e., using previous time estimates, usually in the form of point estimates, as the time priors), which poses the risk that errors in time estimation may easily propagate (Dos Reis et al. 2018). In terms of AOB, previous studies (Gulay et al. 2023; Ward et al. 2021) suggested that Beta-AOB appeared earlier than Gamma-AOB and CAOB, therefore postulating a “Beta-AOB early” hypothesis.

Recent molecular dating analyses employed eukaryotic fossils to calibrate bacterial evolution (Liao et al. 2022; Wang and Luo 2023; Zhang et al. 2023; Wang and Luo 2021). The inclusion of eukaryotic fossils into bacterial dating is justified by the presence of genes observed to have been shared during ancient endosymbiosis events, notably those involving mitochondria and plastids. These eukaryotic fossils provide many more time boundaries, particularly the maxima, for dating analyses (Wang and Luo 2023; Clark and Donoghue 2017). With the incorporation of two eukaryotic fossils each with a maximum constraint, Liao et al. (2022) estimated the origin of anaerobic ammonia oxidation (anammox) bacteria (AnAOB) at ∼2.3 Ga. However, the authors assumed an unrealistically old age of the crown group of eukaryotes at ∼2.3 Ga (Liao et al. 2022), compared with 1.2 - 1.9 Ga reported by most studies (Parfrey et al. 2011; Betts et al. 2018; Wang and Luo 2021). This leaves the possibility that the age of AnAOB was similarly overestimated.

Here, we leverage extensive eukaryotic fossils through plastid endosymbiosis and mitochondrial endosymbiosis to calibrate the evolution of AOB/CAOB and AnAOB. We solve the unrealistically old age problem of the eukaryotes through the Bayesian sequential dating approach (Fig. S5), thereby accurately propagating the timing information available in the eukaryotic portion of the tree to the bacterial nodes. Additionally, we illustrate the impact of using non-vertically transmitted genes on posterior age estimates. Based on a set of predominantly vertically inherited genes, we conclude that AOB predate AnAOB, consistent with the idea that the AOB produce nitrite which serves as the energy source for AnAOB and their electron source for carbon fixation. In contrast to previous studies which reported Beta-AOB as the earliest AOB lineage (Gulay et al. 2023; Ward et al. 2021), our analyses consistently show that Gamma-AOB is the oldest bacterial ammonia oxidizer.

## Results and discussion

### The earliest AOB evolved in Gammaproteobacteria

We first confirmed the phylogenetic distributions of the bacterial ammonia oxidizers and genes. The phylogenomic tree shows that AOB/CAOB are distributed in three deep taxonomic groups (Fig. 1A). The *amoCAB* gene tree (Fig. 1B) confirmed that *amoCAB* arose twice, once in the last common ancestor (LCA) of the Beta-AOB and CAOB and once in the LCA of Gamma-AOB (Soliman and Eldyasti 2018; Lehtovirta-Morley 2018; Khadka et al. 2018). The two versions of *amoCAB* appear to have evolved independently from their paralogs in the CuMMO family (*xmoCAB* genes) (Fig. 1A). Consistent with a previous study (Osborne and Haritos 2018), we found that the *amoCAB* of Gamma-AOB evolved from the *pmoCAB* of methanotrophs, whereas the *amoCAB* of Beta-AOB and CAOB diversified from the *emoCAB* (ethane monooxygenase) and *pxmABC* (ammonia/methane monooxygenase) of non-methanotrophs. To investigate the antiquity of bacterial ammonia oxidizers based on their genome sequences, it is essential to confirm that genome sequences of the basal lineages of this functional group are available. We validated this with the *xmoA* gene tree constructed based on *xmoA* sequences associated with genome sequences and derived from amplicon sequencing of the environmental samples, where we showed that some of the basal lineages of *amo* genes (shaded regions in Fig. 1C) have representatives derived from genome sequences.

**Figure 1.**
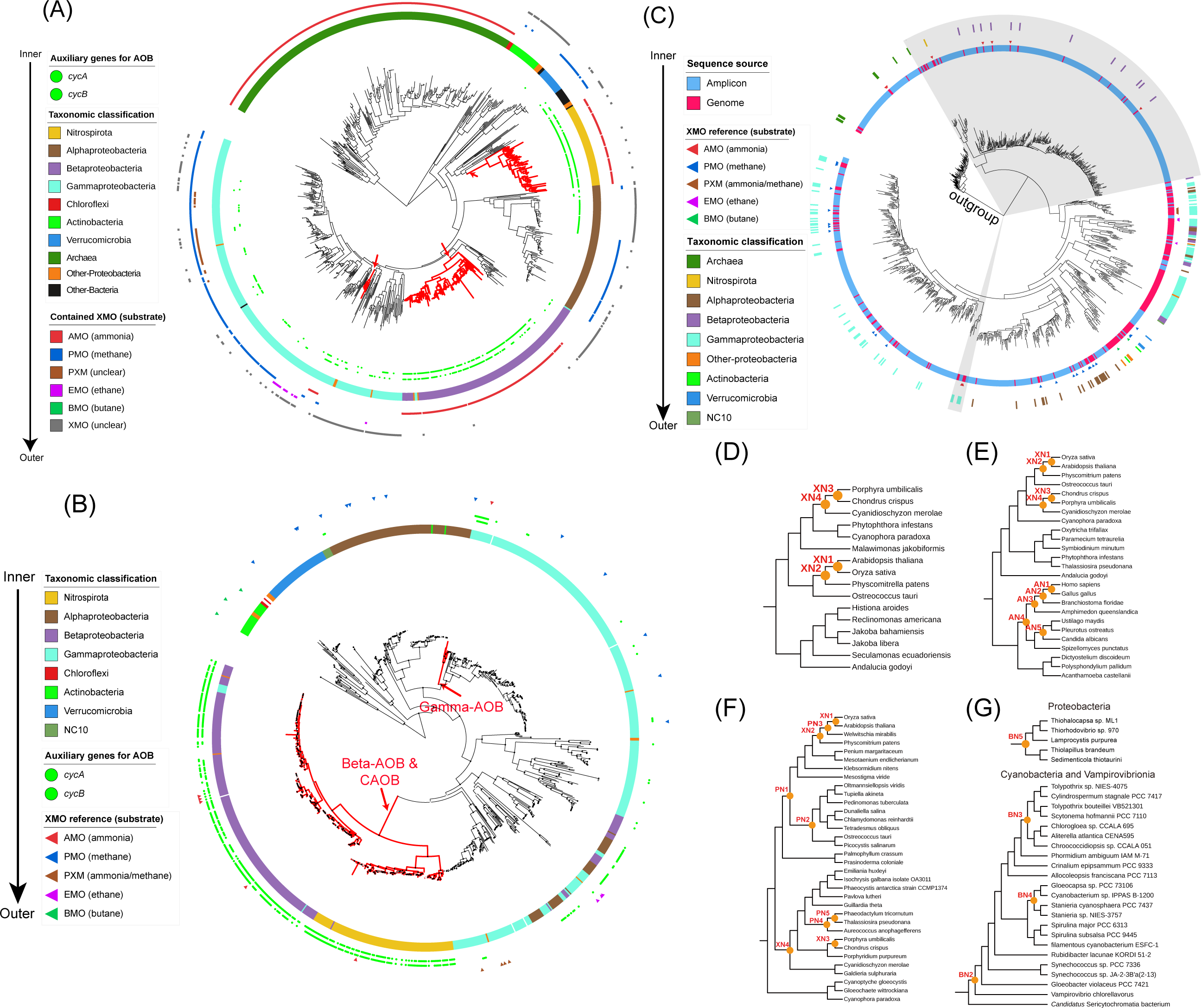
The *xmo* gene phylogeny, the *xmo*-containing genome phylogeny, and the fossil constraints used to calibrate ammonia-oxidizing bacteria (AOB) and complete ammonia oxidizing bacteria (CAOB) evolution. (A) The phylogenomic tree of 848 *xmoA*-containing microbes. The phylogenomic tree was constructed using Phylophlan (see Supplementary Text section 2.3), followed by the outgroup-independent minimum variance (MV) rooting. From inner to outer, the first two rings show the presence of the genes *cycAB*. In the third ring, the color strip represents the major taxonomic lineages derived from the NCBI taxonomy. The remaining six outer rings show the inferred types of XMO according to the concatenated *xmoCAB* gene tree (Fig. 1B). The clades with red branches represent the AOB and CAOB lineages determined based on the presence of *cycAB* and *amoCAB*. (B) The concatenated *xmoCAB* gene tree was generated using a total of 964 *xmoCAB* genes, including 922 identified *xmoCAB* operons and 42 reference sequences (Datasets S1.2 & S1.3) at the amino acid level by IQ-Tree (v1.6.12). The gene tree was rooted with the *bmoCAB* lineage following Khadka et al. (2018). The branches inferred as AOB/CAOB were highlighted in red. From the inner to the outer ring, the first color strip indicates the taxonomic affiliation of the concatenated *xmoCAB*. The circles arranged as two rings represent the presence of the auxiliary genes, *cycAB*, which encode the necessary electron carriers for ammonia oxidation and should be present in the ammonia oxidizers. The outmost ring composed of triangles corresponds to the reference sequences encoding XMO for distinct substrates. (C) The *xmoA* tree was constructed based on the amino acid sequences of *xmoA* homologs obtained from amplicon sequencing and genome sequencing (see Supplementary Text section 2.1). The phylogeny was rooted at the *xmoA* sequences from the archaeal lineages, including *Cenarchaeum*, *Candidatus* Nitrosopumilus, and *Candidatus* Nitrosothermus. From the inner to the outer ring, two shaded regions represent the lineages inferred as the *amoCAB* from AOB/CAOB. The first ring represents the type of the sequence, either amplicon or genome sequence. The next ring composed of triangle corresponds to the reference sequences (see Supplementary Text section 1.1) encoding XMO for distinct substrates. The reference sequences are used to indicate the lineage of XMO with specific substrates. The outermost ring shows the taxonomic affiliation of the tip nodes from genome sequence. (D-G) Calibration nodes for molecular dating analyses using strategies “Mito24” (D), “Gomez19” (E), “Plastid39” (F), and “Battistuzzi25” (G). The eukaryotic topologies (D, E, and F) were pruned from the topologies provided by the previous studies (see Supplementary Text section 3.3), and the bacterial topology (G) was pruned from the tree constructed based on the selected 106 genomes (see Supplementary Text section 2.4). The internal nodes with yellow circles represent the calibration nodes (Datasets S2.1 & S2.2).

We implemented three strategies (“Plastid39”, “Mito24”, and “Gomez19”; the number informs how many genes were used in each strategy), each within a Bayesian sequential dating framework. They all enable the use of fossil-rich eukaryotic lineages to calibrate fossil-poor bacterial lineages by leveraging the plastid endosymbiosis, the mitochondrial endosymbiosis, and the horizontal gene transfer events that relocated some genes from mitochondria to eukaryotic nuclear chromosomes following the mitochondrial endosymbiosis, respectively. Additionally, we implemented a fourth strategy (“Battistuzzi25”) which exclusively uses the 25 conserved genes in bacteria (*Battistuzzi25* in Dataset S1.5) and the fossil calibrations involving Cyanobacteria and purple sulfur bacteria (PSB). Using either 3.5 Ga or 4.5 Ga as the root maximum (Fig. S4), the bacteria-only strategy “Battistuzzi25”, which does not use eukaryotic fossils and Bayesian sequential dating analysis, estimated the age of the crown Gamma-AOB at 2,019 Ma (95% highest posterior density [HPD]: 1,908 – 2,144 Ma) and 2,310 Ma (95% HPD: 2,160 – 2,465 Ma), respectively. However, all three eukaryotes-associated strategies (“Plastid39”, “Mito24”, and “Gomez19”) each gave similar posterior estimates of the crown Gamma-AOB group, with the differences of less than 10, 20, and 90 Ma, respectively. Therefore, the strategy “Battistuzzi25”, which is heavily affected by the age set for the root maximum, is not further discussed.

Our results consistently support Gamma-AOB as the earliest AOB among the three AOB/CAOB groups, supporting the “Gamma-AOB early” hypothesis (Fig. 2). Specifically, the use of the “Plastid39” strategy gave the posterior ages of the crown groups of Gamma-AOB, Beta-AOB, and CAOB to be 1,667 Ma (95% HPD: 1,429 – 1,868 Ma), 785 Ma (95% HPD: 619 – 979 Ma), and 539 Ma (95% HPD: 397 – 696 Ma), respectively. When the “Mito24” strategy was used, the posterior ages of the three lineages became 1,932 Ma (95% HPD: 1,609 – 2,284 Ma), 961 Ma (95% HPD: 662 – 1,304 Ma), and 1,068 Ma (95% HPD: 737 – 1,401 Ma), respectively. Moreover, using Bayesian sequential approaches, the age of crown group eukaryotes shifted to a younger age (1.6 Ga) compared to the unrealistic estimate (2.3 Ga) presented by Liao et al. (2022) who used the same eukaryote-based calibrations. Further, with the “Gomez19” strategy, we inferred the posterior ages of the three lineages to be 1,740 Ma (95% HPD: 1,571 – 1,944 Ma), 974 Ma (95% HPD: 744 – 1,226 Ma), and 1,057 Ma (95% HPD: 802 –1,298 Ma), respectively (Fig. S3; Dataset S2.3). The use of the “Plastid39” strategy consistently led to younger estimates of AOB/CAOB compared to the use of “Mito24” and “Gomez19” strategies. This is likely because the “Plastid39” strategy included eukaryotic fossils within Cyanobacteria, whereas the other two strategies subtend eukaryotic lineages to Alphaproteobacteria. The differences between “Plastid39” and “Mito24”/ “Gomez19” are likely a result of the larger phylogenetic distance between Cyanobacteria and AOB/CAOB compared to the distance between Alphaproteobacteria and AOB/CAOB.

**Figure 2.**
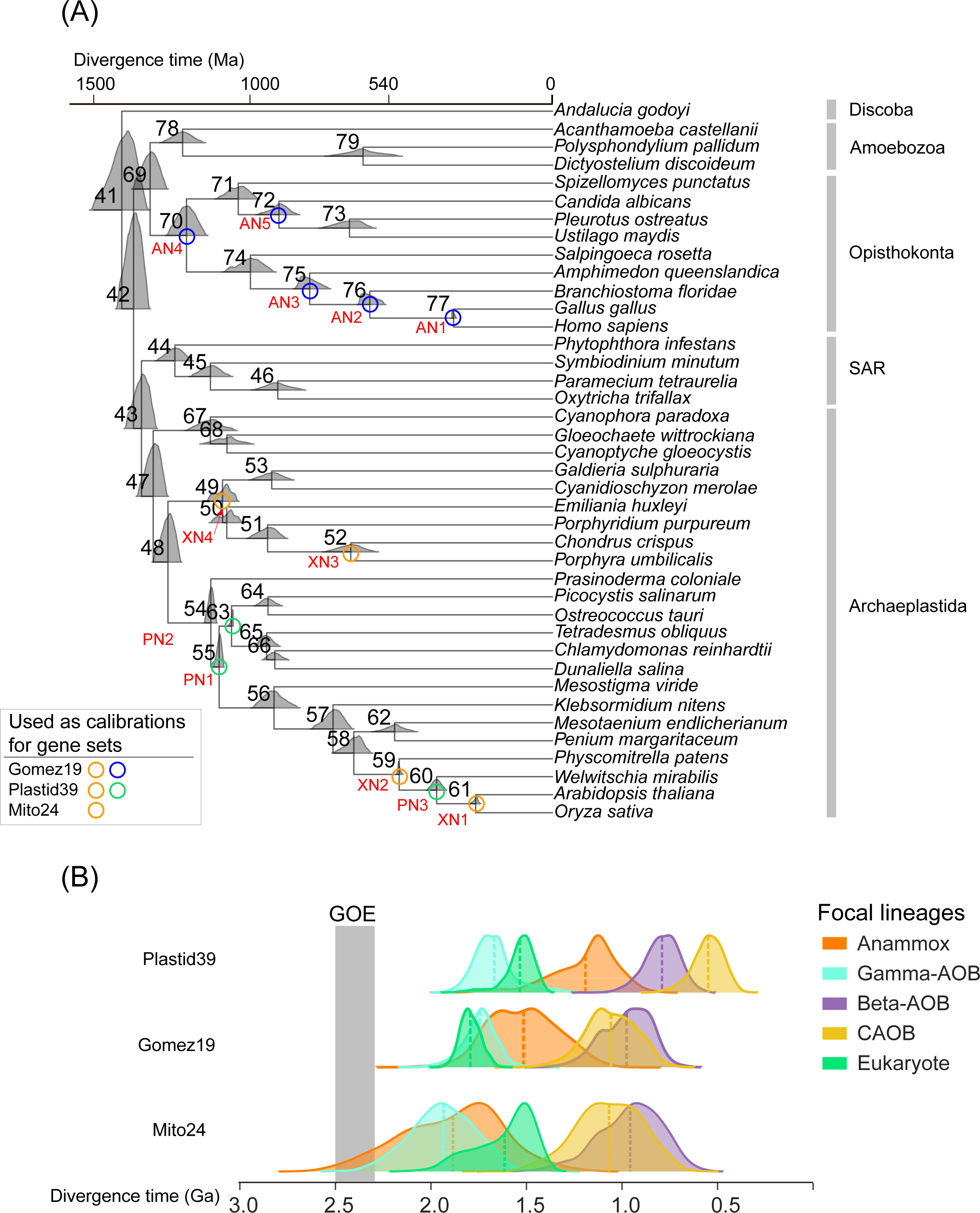
The posterior ages derived from the two-step sequential dating analysis. (A) The posterior ages of the eukaryotic nodes derived from the first-step dating analysis. The posterior ages shown in the density plots at each node were estimated based on 320 orthologs and 12 calibrations (see Supplementary Text section 3.5). The nodes with circles and red calibration node names (XN1 to XN4, PN1 to PN3, and AN1 to AN5) are those with fossil-based calibrations and are those used as calibrations in the second-step dating analysis that includes both eukaryotic and bacterial lineages. The left right panel suggested the used calibrations for each gene set. (B) The posterior ages of five focal lineages derived from the second-step dating analysis. Five focal lineages include Gamma-AOB, Beta-AOB, CAOB, Anammox, and Eukaryote. The estimation of posterior ages utilizing the 12 best-fit distributions (nodes with circles) that were derived from the first-step dating analysis and employed as priors (see Supplementary Text section 3.5). Additionally, the bacterial calibration set B1 was utilized in this process (Dataset S2.1). Three gene sets (*Gomez19*, *Mito24*, and *Plastid39*) and corresponding topologies with eukaryotic genomes as mentioned above (Fig. 1) were employed, respectively. The dashed line within each density plot represents the mean value of each distribution. The full timetrees were shown in the Figure S3.

In contrast to our “Gamma-AOB early” result, previous studies (Gulay et al. 2023; Ward et al. 2021) proposed the “Beta-AOB early” hypothesis. The latter appears to result from inappropriate dating practices. Gulay et al. (2023) used a phylogenomic tree which places the bacterial root at the total group of the candidate phylum Zixibacteria (a member of Fibrobacteres–Chlorobi–Bacteroidetes [FCB] group) for dating analysis. This incorrect root placement led to the grouping of the Cyanobacteria and Proteobacteria, which is very different from the root placement that separates Terrabacteria (including Cyanobacteria) from Gracillicutes (containing Proteobacteria) by modern phylogenetics methods and models (Hug et al. 2016; Coleman et al. 2021; Wang and Luo 2023). Ward et al. (2021) did not include a basal lineage of Gamma-AOB represented by the metagenome-assembled genome (GCA_014859375.1), thus the reported origin of Gamma-AOB at ∼1.0 Ga must be an underestimate. Both studies did not employ the widely used hydrocarbon biomarkers (Okenane and other aromatic carotenoids) to constrain the minimum age of PSB (Chromatiaceae) at 1.6 Ga (Brocks et al. 2005). As PSB is a sister lineage of the Gamma-AOB, including the former and its biomarker-based calibration can help constrain the age of the latter. Without the time constraint based on this biomarker, Ward et al. (2021) estimated the age of the crown group of Gammaproteobacteria at ∼1.3 Ga and Gulay et al. (2023) estimated the age of the crown group of Chromatiaceae at around ∼1.1 Ga, both of which greatly underestimated the origin time of the PSB, and thus probably underestimated the age of AOB.

### Gamma-AOB predate AnAOB

Ammonia oxidation, which produces nitrite, the substrate of AnAOB, is considered as the prerequisite for the origin of AnAOB (Canfield et al. 2010; Liao et al. 2022; Kuypers et al. 2018). In a preliminary analysis, we used the full sets of gene families included in the four strategies. We found that the “Plastid39” and the “Battistuzzi25” strategies support this hypothesis, whereas the “Gomez19” and the “Mito24” strategies instead support the opposite trend where AnAOB predate Gamma-AOB. However, when the gene families that show large ΔLL (Shen et al. 2017; Smith et al. 2020), and thus low congruence with the species tree (Fig. S1), were gradually removed from these datasets, Gamma-AOB shifted towards the ancient whereas AnAOB shifted towards the present (Fig. 3A). Once most of the phylogenetically incongruent genes were removed, Gamma-AOB became older than AnAOB (Fig. 3B).

**Figure 3.**
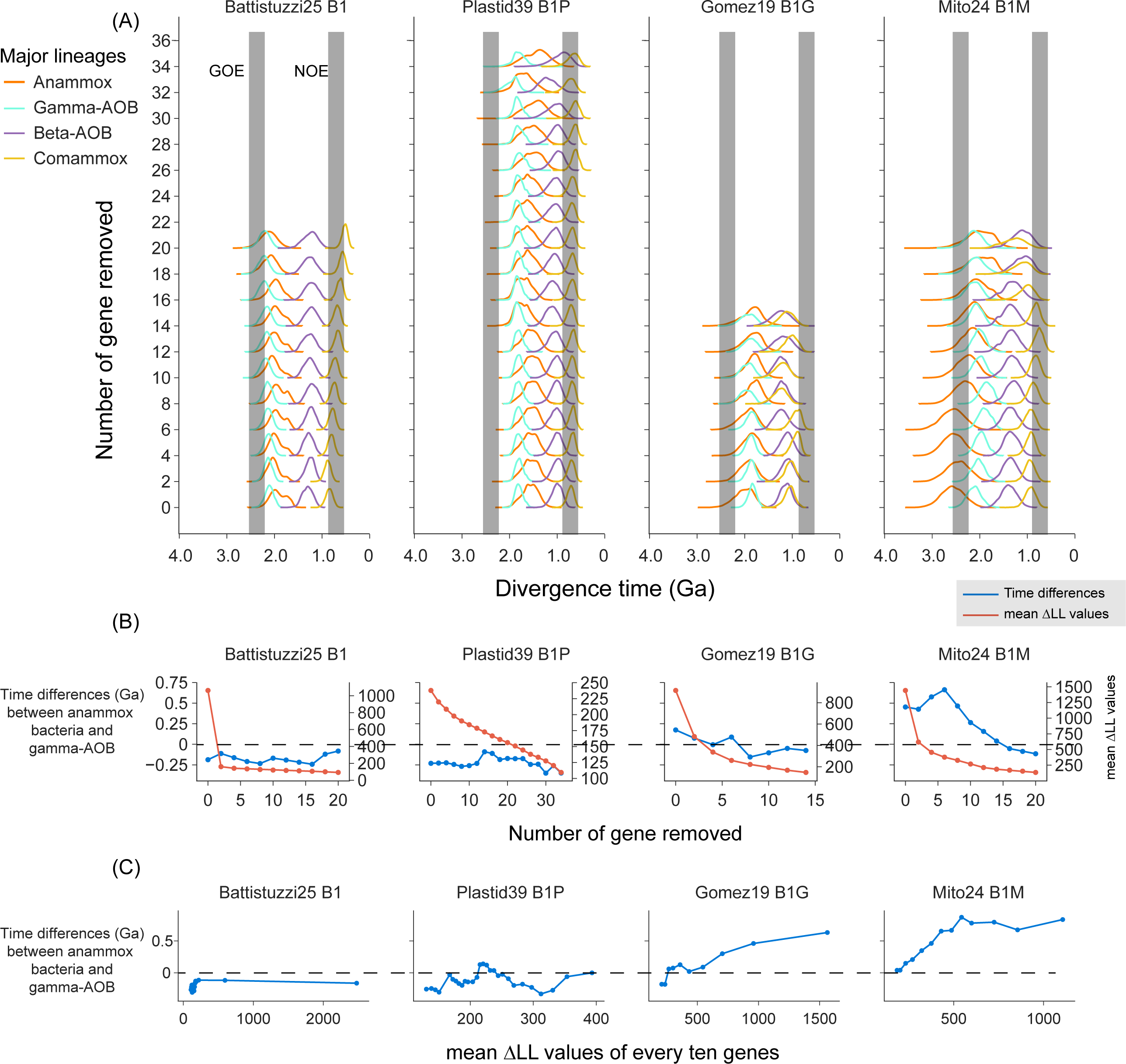
The impact of using phylogenetically incongruent genes on posterior time estimates. For each of the four independent gene sets, the divergence time was estimated using the corresponding calibration set (Dataset S2.1) with MCMCTree (see Supplementary Text Section 3.4). The Bayesian sequential dating approach was not applied for this purpose. The topologies used in each dating analysis (see Supplementary Text Section 3.2) were constructed with bacterial genomes only (“Battistuzzi25”), bacterial genomes and nuclear genomes of eukaryotes (“Gomez19”), bacterial genomes and mitochondrional genomes of eukaryotes (“Mito24”), and bacterial genomes and plastid genomes of eukaryotes (“Plastid39”). (A) The change of posterior age estimates of the gammaproteobacterial AOB (Gamma-AOB), betaproteobacterial AOB (Beta-AOB), comammox bacteria (CAOB), and anammox bacteria (AnAOB) along with the change of the number of genes by gradually removing two genes that are most topologically incongruent with the species tree (i.e., the largest delta log-likelihood [ΔLL]). The time estimates refer to the age of the total group of each lineage. The two vertical grey bars represent the Great Oxidation Event (GOE) occurring at 2,500-2,320 Ma and the Neoproterozoic Oxygenation Event (NOE) at 800-550 Ma, respectively. (B) The change of the time difference between Gamma-AOB and AnAOB along with the change of the number of genes by gradually removing the two genes with the largest ΔLL. The left y-axis (corresponding to the blue curve) shows values calculated by subtracting the mean time estimates of the total group of Gamma-AOB from those of the total group of AnAOB. The right y-axis values (corresponding to the red curve) indicate the mean ΔLL value of the remaining genes following gene removal. (C) The scatter plot shows the estimated age difference between Gamma-AOB and AnAOB of analysis using genes selected with the sliding window approach with a window size of 10 genes and a step size of one gene. The x-axis shows the mean ΔLL of the ten genes selected each time. The y-axis shows values calculated by subtracting the mean time estimates of the total group of Gamma-AOB from the mean time estimates of the total group of AnAOB.

Although this “gene removal” method illustrates the importance of gene verticality on posterior age estimates, it leaves fewer genes for later rounds of molecular dating, and therefore the factor of gene number is not under control. To solve this problem, we implemented a ‘sliding window’ method to shift the gene sets according to their ΔLL values while keeping the number of genes at 10 (Fig. 3C). The results derived from this approach are consistent with those from the “gene removal” approach, supporting the idea that genes with low ΔLL should be used for molecular clock analysis. In sum, our dating analyses with all the four strategies support the “Gamma-AOB early” hypothesis where Gamma-AOB predate AnAOB (Fig. 2B).

### All bacterial ammonia oxidizers postdate the GOE but the origin of ammonia oxidation may be correlated with the GOE

Geochemical studies using N isotopes proposed a Neoarchean origin of biological nitrification (and denitrification) as early as 2.7 Ga (Koehler et al. 2018; Godfrey and Falkowski 2009; Garvin et al. 2009). The isotopic record further suggests that this metabolism became environmentally more widespread around the GOE at 2.3 Ga (Kipp et al. 2018; Zerkle et al. 2017; Garvin et al. 2009; Cheng et al. 2019). The total and crown group of the Gamma-AOB, however, likely evolved slightly later, at 2.1 Ga (95% HPD: 1,864 - 2,375 Ma; “Mito24” strategy) to 1.9 Ga (95% HPD: 1,766 – 1,970 Ma; “Plastid39” strategy) and from 1.9 Ga (95% HPD: 1,609 - 2,284 Ma; “Mito24” strategy) to 1.7 Ga (95% HPD: 1,429 – 1,868 Ma; “Plastid39” strategy), respectively. At that time, nitrification appears to be well established in the sedimentary isotope record (Kump et al. 2011; Stüeken 2013; Godfrey et al. 2013). The timing may also coincide with a later oxygen level peak at 1.9 Ga (Large et al. 2022). Hence, Gamma-AOB, which most likely originated in marine ecosystems, given their habitats today (Lehtovirta-Morley 2018), may have contributed to the isotopic records sourced from the marine environment.

There appears to be a mismatch between the oldest proposed geochemical record of nitrification at 2.7 Ga and the origin of Gamma-AOB at 2.1-1.7 Ga. One possibility is that the geochemical data prior to the GOE has been misinterpreted due to unrecognized alteration or alternative metabolic reactions. Alternatively, the mismatch of the timing between the expression of aerobic N cycling in the geochemical record and the origin of marine bacterial ammonia oxidizers may reveal that microbes other than the traditional Gamma-AOB contributed to ammonia oxidation prior to the GOE. One possibility is the LCA (node X in Fig. S3) shared by Gamma-MOB and Gamma-AOB, which was proposed to possess a dual *xmoCAB* system (two or multiple copies of *xmoCAB*) capable of performing both methane oxidation and ammonia oxidation (Osborne and Haritos 2018). The crown group of the dual *xmoCAB* system-containing bacteria (node X in Fig. S3) was estimated to arise at 1,867 Ma (95% HPD: 1,766 – 1,970 Ma), 1,905 Ma (95% HPD: 1,773 – 2,054 Ma), and 2,108 Ma (95% HPD: 1,864 – 2,375 Ma) using the strategies “Plastid39”, “Gomez19”, and “Mito24”, respectively. Alphaproteobacterial methanotrophs are unlikely to have contributed to the ammonia oxidation before the GOE, as the crown group of Alphaproteobacteria likely occurred at ∼1.9 Ga which was estimated by leveraging many eukaryotic fossils based on the mitochondrial endosymbiosis (Wang and Luo 2021). However, non-proteobacterial methanotrophs, such as those from the Actinobacteria and Verrucomicrobia phyla, lack time estimations for their antiquity. Despite this, they are known to oxidize ammonia and are considered likely contributors to nitrite accumulation preceding the GOE (Klotz and Stein 2008; Schmitz et al. 2021). Intriguingly, the geochemical evidence of nitrification in the Neoarchean occurs in a time interval that is known for unusually light organic carbon isotope values (δ^13^C_org_) that are generally interpreted as evidence of methanotrophy (Eigenbrode and Freeman 2006). This raises the intriguing possibility that the rise of methanotrophs triggered the onset of ammonia oxidation. The increase in atmospheric oxygen levels during the GOE would have removed the atmospheric methane, perhaps causing organisms to favor ammonia over methane as a substrate and resulting in the expansion of ammonia oxidation into other phyla.

Another possible ammonia oxidizer in the Neoarchean is AOA. Ren et al. (2019) estimated the origin of the crown and total group of the AOA at 2.1 Ga (95% HPD: 2,060 – 2,285 Ma) and 2.45 Ga (95% HPD: 2,339 – 2,573 Ma). This age bracket allows for the occurrence of AOA a few hundred million years before the GOE at 2.3 Ga. However, these estimates are vulnerable because i) using secondary calibrations ignores the calibration uncertainty and thereby biasing the posterior age estimates (Dos Reis et al. 2018; Liao et al. 2022); ii) using gene sets without verticality examination results in bias in posterior age estimation, as shown in the present study. Yang et al. (2021) argued that Ren et al. (2019) adopted a strict clock rate model, but in reality Ren et al. (2019) employed an independent rate clock model (see the source code: https://github.com/luolab-cuhk/thaum-dating-project-2019). Although a more careful analysis is required, it remains well possible that the earliest AOA originated at or before the GOE and contributed to the aerobic N cycle at that time. Because AOA originated on the land and remained restricted to terrestrial environments for over a billion years (Ren et al. 2019) and because the sedimentary geochemical records are derived from shallow oceans (Pufahl and Hiatt 2012; Kipp et al. 2018; Zerkle et al. 2017; Garvin et al. 2009), transportation of nitrite from the land to the ocean is required. This interpretation could potentially be reconciled with the rock record, noting that detailed spatial analyses do not exist for these Archean basins, but gradients of nitrate-rich conditions in onshore settings paired with nitrate-depletion offshore have been documented from the mid-Proterozoic (Koehler et al. 2017; Stüeken 2013). Also transport of nitrate from land to sea by rivers has been proposed in models (Thomazo et al. 2018). Such ammonia-oxidizing settings on land are consistent with some evidence of oxidative weathering starting as far back as 2.95 Ga as suggested by chromium (Cr) isotopic data (Crowe et al. 2013; Frei et al. 2016).

### Change of genomic content upon the origin of bacterial ammonia oxidizers

The MDS plot based on the metabolic dissimilarities grouped the nodes representing Gamma-AOB and Beta-AOB together (Fig. 4A). The normalized metabolic dissimilarity between Gamma-AOB and Beta-AOB is smaller than the normalized nucleotide dissimilarity between them (Fig. S6) (*p* < 0.001; Permutation test), suggesting the convergent evolution of metabolic changes from non-AOB to AOB in the two independent origins (beta-AOB and Gamma-AOB). However, the nodes representing CAOB are closer to the non-CAOB Nitrospirota than to the proteobacterial AOB in both MDS plots (Fig. 4A) based on nucleotide dissimilarity. Since CAOB nodes are evolutionarily distant from AOB nodes, but *amoCAB* of CAOB and Beta-AOB form a monophyletic lineage (Fig. 1B), a reasonable evolutionary scenario is that CAOB acquired its *amoCAB* horizontally from Beta-AOB (Palomo et al. 2018) while keeping most of its genomic regions similar to their non-CAOB Nitrospirota counterparts (Palomo et al. 2022).

**Figure 4.**
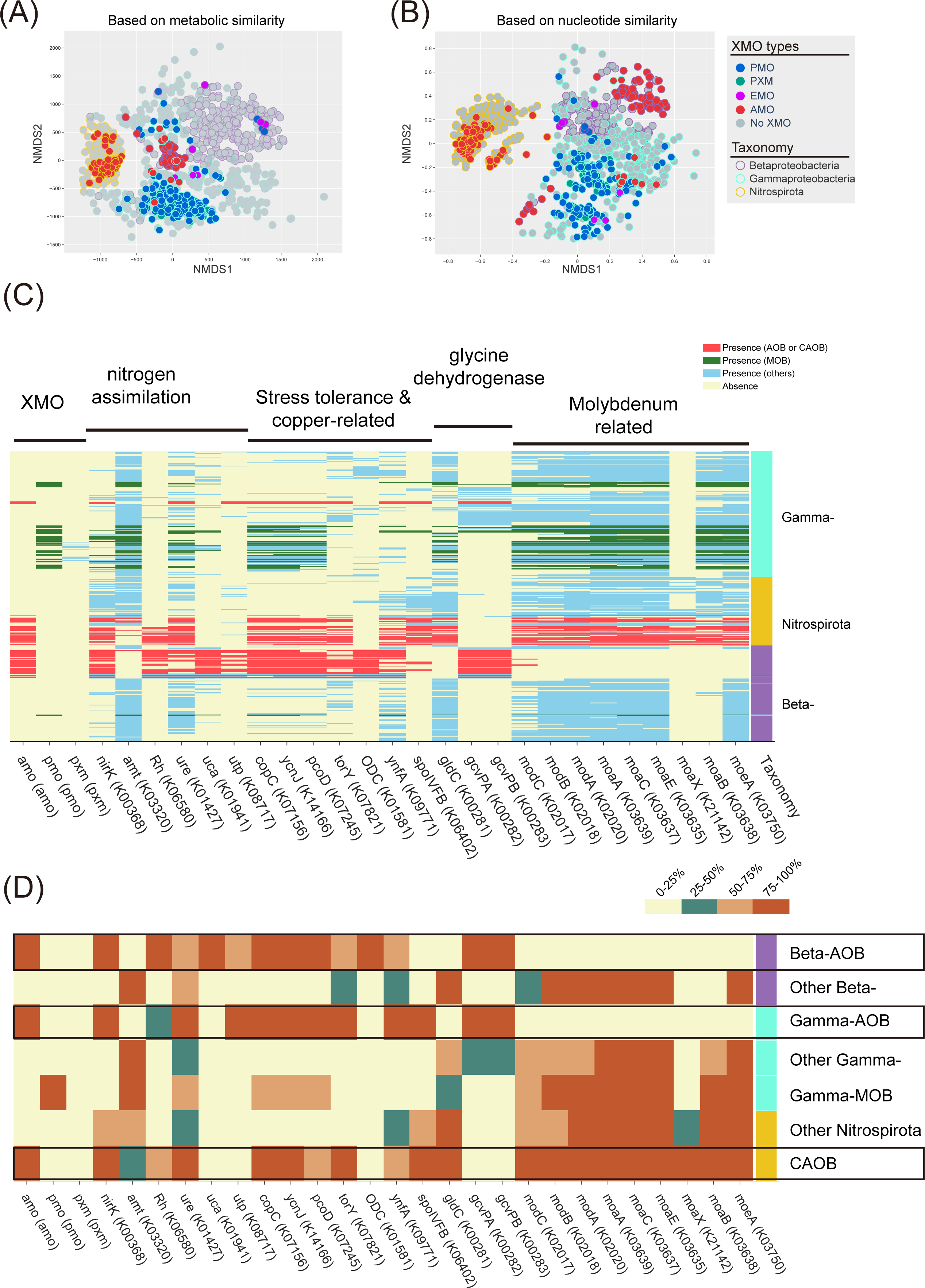
The multidimensional scaling (MDS) analyses of 1,119 genomes including *xmoA*-containing genomes and their phylogenetic relatives that do not contain *xmoA*. The MDS analyses were performed based on metabolic similarity according to the presence and absence of KEGG-annotated genes (A) and nucleotide similarity according to the k-mer based metric (B). Each node represents a genome. The filled color represents the presence and absence of XMO and the type of XMO. The node border color indicates the taxonomic lineages. (C) Distributions of genes statistically associated with *amoCAB* among the 1,119 genomes (see Supplementary Text section 4). The rows and columns represent genomes and KEGG-annotated genes, respectively. To visualize the associations between AMO and other genes, the presence of KEGG-annotated genes in AOB/CAOB, MOB, and other bacteria is highlighted in red, green, and blue, respectively. The absence of the gene is shown in yellow. The rightmost column shows the taxonomic affiliation of the genome. (D) Summary distribution of *amoCAB*-associated genes. The prevalence of the gene among the genomes in a taxonomic lineage or a focal lineage is categorized into four groups distinguished by four colors. Three focal lineages, betaproteobacterial AOB (Beta-AOB), gammaproteobacterial AOB (Gamma-AOB), and complete ammonia-oxidizing bacteria (CAOB), are framed in black boxes. *amo*, ammonia monooxygenase; *pmo*, methane monooxygenase; *pxm*, copper membrane monooxygenase of unknown function; *nirK*, nitrite reductase; *amt*, ammonium transporter; Rh, ammonium transporter; *ure,* urease; *uca*, urea carboxylase; *utp*, urea transporter; *copC*, copper resistance protein C; *ycnJ*, copper transport protein; *torY*, trimethylamine-N-oxide reductase; *ODC*, ornithine decarboxylase; *ynfA*, inner membrane protein; *spoIVFB*, stage IV sporulation protein; *gldC*, homodimeric glycine dehydrogenase; *gcvPAB*, heterotetrameric glycine dehydrogenase; *modABC*, molybdate transport system; *moaABCEX*, molybdenum cofactor biosynthesis; *moeA*, molybdopterin molybdotransferase.

We implemented three association tests to investigate the genomic changes that occurred between non-AOB and AOB as well as between non-CAOB Nitrospirota and CAOB (see Supplementary Text section 4.2). With regard to N-related metabolisms, we found the gains (or presence) of the *nirK* in both Gamma-AOB and Beta-AOB but not in non-AOB close relatives. In contrast, nearly all Nitrospirota (non-CAOB and CAOB) contain *nirK*. The *nirK* gene, which encodes nitrite reductase that is commonly used for reducing nitrite to nitric oxide in denitrification, here is associated with the *amo* gene cluster (Fig. 4C). During ammonia oxidation, the *nirK* nitrite reductase is used to oxidize nitric oxide to nitrite (Caranto and Lancaster 2018; Wijma et al. 2004; Lancaster et al. 2018) and promotes ammonia oxidation at low oxygen levels (5%) (Kozlowski et al. 2014). Therefore, *nirK* likely played an important role in facilitating the emergence of bacterial ammonia oxidizers in early ocean where oxygen levels were very low.

Regarding ammonia transportation and utilization, the *amt* gene for high-affinity ammonia transportation (Khademi et al. 2004) is replaced by the *rh* gene for low-affinity transportation in AOB (Kitano and Saitou 2000; Lupo et al. 2007). Specifically, the majority of non-AOB contain *amt*, while the majority of AOB possess the *rh* gene (for low-affinity transportation) (Fig. 4C). Note that this replacement is also found between non-CAOB and CAOB, which reinforces its importance for bacterial ammonia oxidizers. The primary use of *rh* rather than *amt* in AOB and CAOB (Stein et al. 2007) is likely because the former facilitates bidirectional translocation of ammonia across the cell membrane, which might help adapt to both oligotrophic (importer) and eutrophic (exporter) environments. Furthermore, the *rh* transporter works best in neutral to alkaline conditions (Weidinger et al. 2007; Sogaard et al. 2009) where ammonia, the substrate of aerobic ammonia oxidation, is more available than ammonium (Ullmann et al. 2012).

Both AOB and CAOB can perform ammonia oxidation with urea as the substrate (Koper et al. 2004; Zorz et al. 2018) (Fig. 4C) because they are equipped with metabolic pathways that yield ammonia from urea. This could be either a one-step pathway that uses urease hydrolase encoded by the gene *ure* (Callahan et al. 2005) or a two-step pathway that comprises urea carboxylase and allophanate hydrolase encoded by *uca* and *atzF*, respectively (Kanamori et al. 2004). The *ure* is found in most bacteria and the gene *uca* is uniquely found in Beta-AOB. Note that *atzF* is present in almost all Proteobacteria (Aukema et al. 2020) including the genomes analyzed here (not shown in Fig. 4). Additionally, the gene encoding urea transporter, *utp*, is found in Gamma-AOB and Beta-AOB and missing in CAOB.

Apart from ammonia, copper serves as a main co-factor in AMO and may play an important role in facilitating the origin of bacterial ammonia oxidizers. Marine copper levels sourced from hydrothermal vents and oxidative weathering may have increased during the GOE (Stueeken 2020). The presence of copper in a range of 0.05 to 5 μg/L could significantly increase the rate of ammonia oxidation (Wagner et al. 2016), but higher concentration of copper inhibit the ammonia oxidation (Cela and Sumner 2002). This emphasizes the significance of enzymes in promoting tolerance and preserving equilibrium, such as resistance proteins and transporters. Three copper-related genes show strong correlations including *copC* (copper resistance protein C), *pcoD* (copper resistance protein D), and *ycnJ* (copper transport protein) (Chillappagari et al. 2009). They were found in AOB and CAOB, but not in their non-AOB relatives (Fig. 4). Note that copper also plays an important role in *pmoCAB*, and indeed, these copper-related genes were also found in around 50% of MOB (Fig. 4D). Likewise, molybdenum is the cofactor of nitrite oxidoreductase (NXR) and nitrate reductase (NAR) (Miralles-Robledillo et al. 2019; Schwarz and Mendel 2006), which are involved in the reversible transformation between nitrite and nitrate carried out by NAR and/or NXR. Intriguingly, the genes for molybdenum cofactor biosynthesis (*moaABCXE*, *moeA*) and molybdate transport (*modABC*) are present in CAOB but missing from Gamma-AOB and Beta-AOB (Fig. 4), which likely enable the former to perform the complete ammonia oxidation from ammonia to nitrate. Marine molybdenum levels increased across the GOE and again in the Neoproterozoic (ca. 0.6 Ga; Scott et al. (2008)), which may perhaps be linked to the late origin of CAOB.

### Concluding Remarks

Gamma-AOB are known to have originated in marine environments (Lehtovirta-Morley 2018). Here, we added that they are also the earliest bacterial ammonia oxidizers among the known AOB/CAOB lineages. This finding aligns with the fact that most N isotopic data indicative of nitrification occur in marine sedimentary strata (Kipp et al. 2018; Zerkle et al. 2017). The origin of Gamma-AOB coincides with a time when N isotopes suggest widespread presence of nitrification. Yet, members of the Gamma-AOB crown and total groups, which originated after the GOE (ca. 2.3 Ga), were unlikely to be direct contributors to the earliest isotopic signatures of nitrification, which date as far back as 2.7 Ga. One possibility is that the first ammonia oxidizer was the common ancestor of Gamma-AOB and Gamma-MOB (Osborne and Haritos 2018), given that the latter also has ammonia oxidation activities. Furthermore, other types of methanotrophs, such as those from the Verrucomicrobia and Actinobacteria phyla, have been postulated to possess the capacity for ammonia oxidation (Schmitz et al. 2021; Stein and Klotz 2011). Although the antiquity of these specific methanotrophs were not explored in this study, the specialized extreme environmental niches inhabited by Verrucomicrobia suggest that they could have arisen prior to the GOE. Intriguingly, the carbon isotope record at that time indicates widespread methanotrophy (Eigenbrode & Freeman 2006), opening up the possibility that the origin of methanotrophs went hand in hand with the origin of ammonia oxidation. Another possible contributor is some early AOA. Although AOA originated in terrestrial environments and remained there until well into the mid-Proterozoic (Ren et al. 2019), it is possible that nitrate was generated on land and transported to marine habitats, consistent with evidence of oxidative weathering in the Archean (Crowe et al. 2013; Frei et al. 2016). Moreover, although AOA dominate the ammonia oxidizing communities in the modern ocean (Francis et al. 2005), Gamma-AOB was the only ammonia oxidizer in the earlier ocean before AOA expanded its habitat to the oceans, suggesting a significant role of Gamma-AOB in the early Earth redox cycle.

Of the processes that constitute the N cycle, only nitrification uses oxygen as a substrate (Kuypers et al. 2018). Yet, anaerobic N metabolisms heavily rely on ammonia oxidation to provide nitrite as a substrate to fuel the anaerobic processes. For example, aerobic ammonia oxidation and anammox are usually coupled in modern ecosystems (Liu et al. 2020), which implies their evolutionary order, whereby the former process facilitates the production of nitrite as a substrate for the latter. The conclusion drawn from our analyses supports the idea that Gamma-AOB emerged earlier than AnAOB, which reinforces the hypothesis of a stepwise emergence of these pathways. The ammonia oxidation carried out by methanotrophs prior to the GOE was likely limited to oxygen oases where methane levels may have been low, because in the presence of high methane concentrations may suppress this ammonia oxidation (Bodelier and Laanbroek 2004). The decline of methane after the GOE may thus explain why ammonia oxidation expanded and radiated into other phyla, including Gamma-AOB, and later enable the appearance of AnAOB. Likewise, nitrite oxidation, dissimilatory nitrate reduction to ammonium (DNRA), and denitrification ultimately depend on nitrite produced by aerobic ammonia oxidation. The Proterozoic appearance of Gamma-AOB may therefore explain the expansion of these metabolisms across the tree of life around the same time (Parsons et al. 2021). Our results indicate that nitrate became a more widely available substrate in the Proterozoic, possibly with important consequences for nitrate-assimilating eukaryotes (Anbar and Knoll 2002), the production of N_2_O greenhouse gas (Buick 2007), and the loss of fixed N from the global ocean (Fennel et al. 2008).

## Materials and Methods

The AMO homologs, *xmoCAB* genes, were retrieved by BLASTP using reference protein sequences against the nr database (last updated July 2021). The retrieved 82,848 *xmoA* homologous were clustered by CD-HIT (v4.8.1) (Fu et al. 2012) using 0.9 as the sequence identity threshold then aligned using MAFFT (v7.222) (Katoh and Standley 2013) and trimmed by ClipKit (v1.1.5) (Steenwyk et al. 2020). The protein mixture model (LG+C60) and ultrafast bootstrap test implemented in the IQ-Tree (v1.6.2) (Nguyen et al. 2015) were used to create the gene tree for *xmoA* (Fig. 1C). According to the source of retrieved *xmoA* homologues, we compiled a genome set which include both *xmoCAB*-containing genomes and their closes relatives (7,683 in total; Dataset S1.1) (see Supplementary Texts section 1.2) via Entrez (Kans 2020) and a custom python script (see Data and Code availability). Moreover, the auxiliary genes *hao*, *cycAB*, and *nxrAB* were annotated by using BLASTP (E-value < 1e-50) based on the reference sequences. The phylogenomic tree (Fig. 1B) comprising 848 *xmoA*-containing genomes (Dataset S1.3) was generated by PhyloPhlAn (v3.0) (Segata et al. 2013) using ‘high diversity’ mode. We then identified the genomes containing at least two genes of the *xmoCAB* gene cluster using a custom python script and concatenated the *xmoCAB* alignments for phylogeny reconstruction using IQ-tree under the auto-determined substitution model (LG+C50+G) with 1,000 ultrafast bootstrap replicates. The last common ancestors (LCA) of the three groups of ammonia oxidizers (i.e., Gamma-AOB, Beta-AOB, and CAOB) are recognized based on the presence and absence of the *cycAB* (Fig. 1B) and *xmoCAB* genes.

Due to the large computational cost of Bayesian-based molecular dating analysis, we performed a two-step taxon sampling strategy (see Supplementary Texts section 2.4) to select a total of 106 bacterial genomes and 75 eukaryotic genomes for our dating analysis. For Bayesian dating analysis, four gene sets were compiled including, i) “Battistuzzi25”: 25 universally conserved genes among bacteria (Battistuzzi and Hedges 2009), ii) “Mito24”: 24 conserved genes encoded by mitochondrial genomes (Wang and Luo 2021), iii) “Gomez19”: 19 mitochondria-originated genes that have likely been transferred to the eukaryotic nuclear genome originally identified by (Munoz-Gomez et al. 2022) and that are conserved across the bacterial tree of life (Wang and Luo 2023), and iv) “Plastid39”: 39 genes that are conserved among eukaryotic plastids, cyanobacteria, and our genome sets (Ponce-Toledo et al. 2017). Note that 39 genes were selected out of 79 genes that are conserved among eukaryotic plastids and cyanobacteria based on the distributions of the genes in our genome sets. We searched these marker genes in our genome sets by using HMMER (v3.2.1) (Mistry et al. 2013) or BLASTP 2.9.0 (Johnson et al. 2008) with the E-value threshold at 1e-20 (see Supplementary Texts section 3.1). The presence and absence of these marker genes are shown in Dataset S1.8.

The molecular dating analysis was performed by using the program MCMCTree in the PAML package (4.9j) (Yang 2007). The species tree topology used in molecular dating analysis was first reconstructed with 106 bacterial genomes using the LG+C60+F+G model with IQ-Tree and was then expanded by incorporating representative species that used in in previous studies (see Supplementary Text section 3.3) using ETE (Huerta-Cepas et al. 2016) (Figure 1D to Figure G). To select the best-fit clock model, we conducted an approximate likelihood method (Dos Reis and Z 2011) by using MCMCTree and calculated the Bayes Factor of the independent rate model and the auto-correlated rate model. In order to evaluate the impact of gene family verticality on posterior time estimates, we employed the “gene removal” and the “sliding window” method to compile a series of subset of molecular data based on the delta log likelihood (ΔLL) values of the gene families in each strategy (see Supplementary Text section 3.4). The gene-removal approach step-wisely removes genes with larger ΔLL, while the sliding window approach uses every ten genes following the sorting of all genes according to their ΔLL values. Then, we employed Bayesian sequential strategy (Dos Reis et al. 2012) to refine the bacterial timeline estimation using the gene sets “Plastid39”, “Gomez19” and “Mito24” (see Supplementary Text section 3.5). We perform the first-step dating analysis using 40 eukaryotic nuclear genomes (Dataset S1.7), 320 orthologues (Strassert et al. 2021), and 14 calibrations (XN1 to XN4, PN1 to PN3, and AN1 to AN5 in Dataset S2.1). The estimated posterior ages were subsequently used as the time priors for the second-step dating analysis for “Plastid39”, “Gomez19” and “Mito24”, respectively. The convergence of the molecular dating analysis was evaluated by comparing the posterior dates of two independent runs.

To identify the metabolic differences between AOB and non-AOB and between CAOB and non-CAOB, we sampled 1149 genomes from GenBank (Supplementary Text section 4.1). We then assessed the dissimilarities between genomes using non-metric multidimensional scaling (MDS) algorithm, which quantifies the metabolic dissimilarity and the nucleotide dissimilarity by using the pairwise Manhattan distance matrix and the pairwise mash distance matrix, respectively (Ondov et al. 2016). The protein-coding genes of the sampled genomes were annotated by using KEGG (Aramaki et al. 2020) (see Supplemental Texts section 4.1) and then subjected to Fisher-exact test, Pearson correlation test, and the phylogenetic signal test (phylosig) (Blomberg et al. 2003; Revell 2012) to identify their associations with *amoCAB*. For each gene, the tests were made between i) AOB, CAOB, and others, ii) AOB and non-AOB proteobacteria, iii) Gamma-AOB and non-AOB gammaproteobacteria, iv) Beta-AOB and non-AOB betaproteobacteria, and v) CAOB and non-CAOB Nitrospirota. The resulting *p*-values were corrected using the Benjamini-Hochberg false discovery rate (FDR) procedure. Finally, genes with corrected *P*-values less than 0.05 in five groups of comparisons are manually examined according to the correlation coefficient and the phylogenetic signal derived from the R package “phylosig” (Revell 2012).

## Acknowledgements

We thank Weizhi Song for his help in editing an earlier version of the manuscript. This work is supported by the Hong Kong Research Grants Council (RGC) General Research Fund (GRF) (14107823), the Natural Science Foundation of China (42293294), and the Hong Kong Research Grants Council Area of Excellence Scheme (AoE/M-403/16).

## Data and Code availability

The input and output files of our dating analysis and the custom scripts are deposited in the online repository https://github.com/luolab-cuhk/AOB-origin.

## References

Anbar, Ariel D, and Andrew H Knoll. 2002. ‘Proterozoic ocean chemistry and evolution: a bioinorganic bridge?’, science, 297: 1137–42.

Aramaki, T., R. Blanc-Mathieu, H. Endo, K. Ohkubo, M. Kanehisa, S. Goto, and H. Ogata. 2020. ‘KofamKOALA: KEGG Ortholog assignment based on profile HMM and adaptive score threshold’, Bioinformatics, 36: 2251–52.

Aukema, K. G., L. J. Tassoulas, S. L. Robinson, J. F. Konopatski, M. D. Bygd, and L. P. Wackett. 2020. ‘Cyanuric Acid Biodegradation via Biuret: Physiology, Taxonomy, and Geospatial Distribution’, Appl Environ Microbiol, 86.

Battistuzzi, F. U., and S. B. Hedges. 2009. ‘A major clade of prokaryotes with ancient adaptations to life on land’, Mol Biol Evol, 26: 335–43.

Betts, H. C., M. N. Puttick, J. W. Clark, T. A. Williams, P. C. J. Donoghue, and D. Pisani. 2018. ‘Integrated genomic and fossil evidence illuminates life’s early evolution and eukaryote origin’, Nat Ecol Evol, 2: 1556–62.

Blomberg, Simon P, Theodore Garland Jr, and Anthony R Ives. 2003. ‘Testing for phylogenetic signal in comparative data: behavioral traits are more labile’, Evolution, 57: 717–45.

Bodelier, Paul LE, and Hendrikus J Laanbroek. 2004. ‘Nitrogen as a regulatory factor of methane oxidation in soils and sediments’, FEMS microbiology ecology, 47: 265–77.

Brocks, J. J., G. D. Love, R. E. Summons, A. H. Knoll, G. A. Logan, and S. A. Bowden. 2005. ‘Biomarker evidence for green and purple sulphur bacteria in a stratified Palaeoproterozoic sea’, Nature, 437: 866–70.

Buick, R. 2007. ‘Did the Proterozoic ‘Canfield Ocean’ cause a laughing gas greenhouse?’, Geobiology, 5: 97–100.

Callahan, B. P., Y. Yuan, and R. Wolfenden. 2005. ‘The burden borne by urease’, J Am Chem Soc, 127: 10828–9.

Canfield, D. E., A. N. Glazer, and P. G. Falkowski. 2010. ‘The evolution and future of Earth’s nitrogen cycle’, Science, 330: 192–6.

Caranto, J. D., and K. M. Lancaster. 2018. ‘Correction for Caranto and Lancaster, Nitric oxide is an obligate bacterial nitrification intermediate produced by hydroxylamine oxidoreductase’, Proc Natl Acad Sci USA, 115: E8325.

Cela, Shkelqim, and Malcolm E Sumner. 2002. ‘Critical concentrations of copper, nickel, lead, and cadmium in soils based on nitrification’, Communications in Soil Science and Plant Analysis, 33: 19–30.

Cheng, Chen, Vincent Busigny, Magali Ader, Christophe Thomazo, Carine Chaduteau, and Pascal Philippot. 2019. ‘Nitrogen isotope evidence for stepwise oxygenation of the ocean during the Great Oxidation Event’, Geochimica et Cosmochimica Acta, 261: 224–47.

Chillappagari, S., M. Miethke, H. Trip, O. P. Kuipers, and M. A. Marahiel. 2009. ‘Copper Acquisition Is Mediated by YcnJ and Regulated by YcnK and CsoR in Bacillus subtilis’, Journal of Bacteriology, 191: 2362–70.

Clark, J. W., and P. C. J. Donoghue. 2017. ‘Constraining the timing of whole genome duplication in plant evolutionary history’, Proc Biol Sci, 284.

Coleman, G. A., A. A. Davin, T. A. Mahendrarajah, L. L. Szantho, A. Spang, P. Hugenholtz, G. J. Szollosi, and T. A. Williams. 2021. ‘A rooted phylogeny resolves early bacterial evolution’, Science, 372.

Crowe, S. A., L. N. Dossing, N. J. Beukes, M. Bau, S. J. Kruger, R. Frei, and D. E. Canfield. 2013. ‘Atmospheric oxygenation three billion years ago’, Nature, 501: 535–8.

Dos Reis, M, Gunnell G. F, Barba-Montoya J, Wilkins A, Yang Z, and Yoder A D. 2018. ‘Using phylogenomic data to explore the effects of relaxed clocks and calibration strategies on divergence time estimation: primates as a test case’, Systematic Biology, 67: 594–615.

Dos Reis, M, Inoue J, Hasegawa M, Asher R. J., Donoghue PCJ, and Yang Z. 2012. ‘Phylogenomic datasets provide both precision and accuracy in estimating the timescale of placental mammal phylogeny’, Proc Biol Sci, 279: 3491–500.

Dos Reis, M, Donoghue PCJ, and Yang Z. 2016. ‘Bayesian molecular clock dating of species divergences in the genomics era’, Nat Rev Genet, 17: 71–80.

Dos Reis, M, and Yang Z. 2011. ‘Approximate likelihood calculation on a phylogeny for Bayesian estimation of divergence times’, Mol Biol Evol, 28: 2161–72.

Eigenbrode, Jennifer L, and Katherine H Freeman. 2006. ‘Late Archean rise of aerobic microbial ecosystems’, Proc Natl Acad Sci USA, 103: 15759–64.

Fennel, Katja, Mick Follows, and Paul G Falkowski. 2005. ‘The co-evolution of the nitrogen, carbon and oxygen cycles in the Proterozoic ocean’, American Journal of Science, 305: 526–45.

Fennel, Katja, John Wilkin, Michael Previdi, and Raymond Najjar. 2008. ‘Denitrification effects on air-sea CO2 flux in the coastal ocean: Simulations for the northwest North Atlantic’, Geophysical Research Letters, 35.

Francis, C. A., K. J. Roberts, J. M. Beman, A. E. Santoro, and B. B. Oakley. 2005. ‘Ubiquity and diversity of ammonia-oxidizing archaea in water columns and sediments of the ocean’, Proc Natl Acad Sci USA, 102: 14683–8.

Frei, R., S. A. Crowe, M. Bau, A. Polat, D. A. Fowle, and L. N. Dossing. 2016. ‘Oxidative elemental cycling under the low O2 Eoarchean atmosphere’, Sci Rep, 6: 21058.

Fu, L. M., B. F. Niu, Z. W. Zhu, S. T. Wu, and W. Z. Li. 2012. ‘CD-HIT: accelerated for clustering the next-generation sequencing data’, Bioinformatics, 28: 3150–52.

Garvin, J., R. Buick, A. D. Anbar, G. L. Arnold, and A. J. Kaufman. 2009. ‘Isotopic Evidence for an Aerobic Nitrogen Cycle in the Latest Archean’, Science, 323: 1045–48.

Godfrey, L. V., and P. G. Falkowski. 2009. ‘The cycling and redox state of nitrogen in the Archaean ocean’, Nature Geoscience, 2: 725–29.

Godfrey, Linda V, Simon W Poulton, Gray E Bebout, and Philip W Fralick. 2013. ‘Stability of the nitrogen cycle during development of sulfidic water in the redox-stratified late Paleoproterozoic Ocean’, Geology, 41: 655–58.

Gulay, A., G. Fournier, B. F. Smets, and P. R. Girguis. 2023. ‘Proterozoic Acquisition of Archaeal Genes for Extracellular Electron Transfer: A Metabolic Adaptation of Aerobic Ammonia-Oxidizing Bacteria to Oxygen Limitation’, Mol Biol Evol, 40: msad161.

Huerta-Cepas, J., F. Serra, and P. Bork. 2016. ‘ETE 3: Reconstruction, Analysis, and Visualization of Phylogenomic Data’, Mol Biol Evol, 33: 1635–8.

Hug, L. A., B. J. Baker, K. Anantharaman, C. T. Brown, A. J. Probst, C. J. Castelle, C. N. Butterfield, A. W. Hernsdorf, Y. Amano, K. Ise, Y. Suzuki, N. Dudek, D. A. Relman, K. M. Finstad, R. Amundson, B. C. Thomas, and J. F. Banfield. 2016. ‘A new view of the tree of life’, Nat Microbiol, 1: 16048.

Johnson, M., I. Zaretskaya, Y. Raytselis, Y. Merezhuk, S. McGinnis, and T. L. Madden. 2008. ‘NCBI BLAST: a better web interface’, Nucleic Acids Res, 36: W5–9.

Jones, R. D., and R. Y. Morita. 1983. ‘Methane Oxidation by Nitrosococcus oceanus and Nitrosomonas europaea’, Appl Environ Microbiol, 45: 401–10.

Kanamori, T., N. Kanou, H. Atomi, and T. Imanaka. 2004. ‘Enzymatic characterization of a prokaryotic urea carboxylase’, Journal of Bacteriology, 186: 2532–39.

Kans, Jonathan. 2020. ‘Entrez direct: E-utilities on the UNIX command line.’ in, Entrez Programming Utilities Help [Internet] (National Center for Biotechnology Information (US)).

Katoh, K., and D. M. Standley. 2013. ‘MAFFT multiple sequence alignment software version 7: improvements in performance and usability’, Mol Biol Evol, 30: 772–80.

Khademi, Shahram, Joseph O’Connell III, Jonathan Remis, Yaneth Robles-Colmenares, Larry JW Miercke, and Robert M Stroud. 2004. ‘Mechanism of ammonia transport by Amt/MEP/Rh: structure of AmtB at 1.35 A’, Science, 305: 1587–94.

Khadka, R., L. Clothier, L. Wang, C. K. Lim, M. G. Klotz, and P. F. Dunfield. 2018. ‘Evolutionary History of Copper Membrane Monooxygenases’, Front Microbiol, 9: 2493.

Kipp, Michael A, Eva E Stüeken, Misuk Yun, Andrey Bekker, and Roger Buick. 2018. ‘Pervasive aerobic nitrogen cycling in the surface ocean across the Paleoproterozoic Era’, Earth and Planetary Science Letters, 500: 117–26.

Kitano, T., and N. Saitou. 2000. ‘Evolutionary history of the Rh blood group-related genes in vertebrates’, Immunogenetics, 51: 856–62.

Klotz, M. G., and L. Y. Stein. 2008. ‘Nitrifier genomics and evolution of the nitrogen cycle’, FEMS Microbiol Lett, 278: 146–56.

Koehler, Matthew C, Roger Buick, Michael A Kipp, Eva E Stüeken, and Jonathan Zaloumis. 2018. ‘Transient surface ocean oxygenation recorded in the∼ 2.66-Ga Jeerinah Formation, Australia’, Proc Natl Acad Sci USA, 115: 7711–16.

Koehler, Matthew C, Eva E Stüeken, Michael A Kipp, Roger Buick, and Andrew H Knoll. 2017. ‘Spatial and temporal trends in Precambrian nitrogen cycling: A Mesoproterozoic offshore nitrate minimum’, Geochimica et Cosmochimica Acta, 198: 315–37.

Koper, T. E., A. F. El-Sheikh, J. M. Norton, and M. G. Klotz. 2004. ‘Urease-encoding genes in ammonia-oxidizing bacteria’, Appl Environ Microbiol, 70: 2342–8.

Kozlowski, J. A., J. Price, and L. Y. Stein. 2014. ‘Revision of N2O-producing pathways in the ammonia-oxidizing bacterium Nitrosomonas europaea ATCC 19718’, Appl Environ Microbiol, 80: 4930–5.

Kump, L. R., C. Junium, M. A. Arthur, A. Brasier, A. Fallick, V. Melezhik, A. Lepland, A. E. Crne, and G. Luo. 2011. ‘Isotopic evidence for massive oxidation of organic matter following the great oxidation event’, Science, 334: 1694–6.

Kuypers, M. M. M., H. K. Marchant, and B. Kartal. 2018. ‘The microbial nitrogen-cycling network’, Nat Rev Microbiol, 16: 263–76.

Lancaster, K. M., J. D. Caranto, S. H. Majer, and M. A. Smith. 2018. ‘Alternative Bioenergy: Updates to and Challenges in Nitrification Metalloenzymology’, Joule, 2: 421–41.

Large, Ross R., Robert M. Hazen, Shaunna M. Morrison, Dan D. Gregory, Jeffrey A. Steadman, and Indrani Mukherjee. 2022. ‘Evidence that the GOE was a prolonged event with a peak around 1900 Ma’, Geosystems and Geoenvironment, 1: 100036.

Lehtovirta-Morley, L. E. 2018. ‘Ammonia oxidation: Ecology, physiology, biochemistry and why they must all come together’, Fems Microbiology Letters, 365.

Liao, T., S. Wang, E. E. Stueken, and H. Luo. 2022. ‘Phylogenomic Evidence for the Origin of Obligate Anaerobic Anammox Bacteria Around the Great Oxidation Event’, Mol Biol Evol, 39.

Liu, C., L. Hou, M. Liu, Y. Zheng, G. Yin, H. Dong, X. Liang, X. Li, D. Gao, and Z. Zhang. 2020. ‘In situ nitrogen removal processes in intertidal wetlands of the Yangtze Estuary’, J Environ Sci (China), 93: 91–97.

Lupo, D., X. D. Li, A. Durand, T. Tomizaki, B. Cherif-Zahar, G. Matassi, M. Merrick, and F. K. Winkler. 2007. ‘The 1.3-angstrom resolution structure of Nitrosomonas europaea Rh50 and mechanistic implications for NH3 transport by Rhesus family proteins’, Proc Natl Acad Sci USA, 104: 19303–08.

Miralles-Robledillo, J. M., J. Torregrosa-Crespo, R. M. Martinez-Espinosa, and C. Pire. 2019. ‘DMSO Reductase Family: Phylogenetics and Applications of Extremophiles’, Int J Mol Sci, 20.

Mistry, J., R. D. Finn, S. R. Eddy, A. Bateman, and M. Punta. 2013. ‘Challenges in homology search: HMMER3 and convergent evolution of coiled-coil regions’, Nucleic Acids Res, 41: e121.

Munoz-Gomez, S. A., E. Susko, K. Williamson, L. Eme, C. H. Slamovits, D. Moreira, P. Lopez-Garcia, and A. J. Roger. 2022. ‘Site-and-branch-heterogeneous analyses of an expanded dataset favour mitochondria as sister to known Alphaproteobacteria’, Nat Ecol Evol, 6: 253–62.

Nguyen, L. T., H. A. Schmidt, A. von Haeseler, and B. Q. Minh. 2015. ‘IQ-TREE: a fast and effective stochastic algorithm for estimating maximum-likelihood phylogenies’, Mol Biol Evol, 32: 268–74.

O’neill, JG, and JF Wilkinson. 1977. ‘Oxidation of ammonia by methane-oxidizing bacteria and the effects of ammonia on methane oxidation’, Microbiology, 100: 407–12.

Ondov, B. D., T. J. Treangen, P. Melsted, A. B. Mallonee, N. H. Bergman, S. Koren, and A. M. Phillippy. 2016. ‘Mash: fast genome and metagenome distance estimation using MinHash’, Genome Biol, 17: 132.

Osborne, C. D., and V. S. Haritos. 2018. ‘Horizontal gene transfer of three co-inherited methane monooxygenase systems gave rise to methanotrophy in the Proteobacteria’, Mol Phylogenet Evol, 129: 171–81.

Ossa, Frantz Ossa, Jorge E Spangenberg, Andrey Bekker, Stephan König, Eva E Stüeken, Axel Hofmann, Simon W Poulton, Aierken Yierpan, Maria I Varas-Reus, and Benjamin Eickmann. 2022. ‘Moderate levels of oxygenation during the late stage of Earth’s Great Oxidation Event’, Earth Planet Sci Lett, 594: 117716.

Pajares, Silvia, and Ramiro Ramos. 2019. ‘Processes and microorganisms involved in the marine nitrogen cycle: knowledge and gaps’, Frontiers in Marine Science, 6: 739.

Palomo, A., A. Dechesne, O. X. Cordero, and B. F. Smets. 2022. ‘Evolutionary Ecology of Natural Comammox Nitrospira Populations’, mSystems, 7: e0113921.

Palomo, A., A. G. Pedersen, S. J. Fowler, A. Dechesne, T. Sicheritz-Ponten, and B. F. Smets. 2018. ‘Comparative genomics sheds light on niche differentiation and the evolutionary history of comammox Nitrospira’, ISME J, 12: 1779–93.

Parfrey, L. W., D. J. Lahr, A. H. Knoll, and L. A. Katz. 2011. ‘Estimating the timing of early eukaryotic diversification with multigene molecular clocks’, Proc Natl Acad Sci USA, 108: 13624–9.

Park, S. J., A. S. Andrei, P. A. Bulzu, V. S. Kavagutti, R. Ghai, and A. C. Mosier. 2020. ‘Expanded Diversity and Metabolic Versatility of Marine Nitrite-Oxidizing Bacteria Revealed by Cultivation- and Genomics-Based Approaches’, Appl Environ Microbiol, 86.

Parsons, Chris, Eva E Stüeken, Caleb J Rosen, Katherine Mateos, and Rika E Anderson. 2021. ‘Radiation of nitrogen-metabolizing enzymes across the tree of life tracks environmental transitions in Earth history’, Geobiology, 19: 18–34.

Pellerin, Alice, Christophe Thomazo, Magali Ader, Johanna Marin-Carbonne, Julien Alleon, Emmanuelle Vennin, and Axel Hofmann. 2023. "Iron-mediated anaerobic ammonium oxidation recorded in the early Archean ferruginous ocean." In.: Wiley Online Library.

Ponce-Toledo, Rafael I, Philippe Deschamps, Purificación López-García, Yvan Zivanovic, Karim Benzerara, and David Moreira. 2017. ‘An early-branching freshwater cyanobacterium at the origin of plastids’, Current Biology, 27: 386–91.

Preena, P. G., V. J. R. Kumar, and I. S. B. Singh. 2021. ‘Nitrification and denitrification in recirculating aquaculture systems: the processes and players’, Reviews in Aquaculture, 13: 2053–75.

Pufahl, PK, and EE Hiatt. 2012. ‘Oxygenation of the Earth’s atmosphere–ocean system: a review of physical and chemical sedimentologic responses’, Marine and Petroleum Geology, 32: 1–20.

Ren, M., X. Feng, Y. Huang, H. Wang, Z. Hu, S. Clingenpeel, B. K. Swan, M. M. Fonseca, D. Posada, R. Stepanauskas, J. T. Hollibaugh, P. G. Foster, T. Woyke, and H. Luo. 2019. ‘Phylogenomics suggests oxygen availability as a driving force in Thaumarchaeota evolution’, ISME J, 13: 2150–61.

Revell, Liam J. 2012. ‘phytools: an R package for phylogenetic comparative biology (and other things)’, Methods in ecology and evolution: 217–23.

Robinson, Rebecca S, Markus Kienast, Ana Luiza Albuquerque, Mark Altabet, Sergio Contreras, Ricardo De Pol Holz, Nathalie Dubois, Roger Francois, Eric Galbraith, and Ting-Chang Hsu. 2012. ‘A review of nitrogen isotopic alteration in marine sediments’, Paleoceanography, 27.

Schmitz, R. A., S. H. Peeters, W. Versantvoort, N. Picone, A. Pol, M. S. M. Jetten, and H. J. M. Op den Camp. 2021. ‘Verrucomicrobial methanotrophs: ecophysiology of metabolically versatile acidophiles’, FEMS Microbiol Rev, 45.

Schwarz, G., and R. R. Mendel. 2006. ‘Molybdenum cofactor biosynthesis and molybdenum enzymes’, Annu Rev Plant Biol, 57: 623–47.

Scott, C, TW Lyons, A Bekker, Yan-an Shen, SW Poulton, Xue-lei Chu, and AD Anbar. 2008. ‘Tracing the stepwise oxygenation of the Proterozoic ocean’, Nature, 452: 456–59.

Segata, N., D. Bornigen, X. C. Morgan, and C. Huttenhower. 2013. ‘PhyloPhlAn is a new method for improved phylogenetic and taxonomic placement of microbes’, Nat Commun, 4: 2304.

Shen, Xing-Xing, Chris Todd Hittinger, and Antonis Rokas. 2017. ‘Contentious relationships in phylogenomic studies can be driven by a handful of genes’, Nat Ecol Evol, 1: 0126.

Simon, J., and M. G. Klotz. 2013. ‘Diversity and evolution of bioenergetic systems involved in microbial nitrogen compound transformations’, Biochimica Et Biophysica Acta-Bioenergetics, 1827: 114–35.

Smith, S. A., N. Walker-Hale, J. F. Walker, and J. W. Brown. 2020. ‘Phylogenetic Conflicts, Combinability, and Deep Phylogenomics in Plants’, Syst Biol, 69: 579–92.

Sogaard, R., M. Alsterfjord, N. Macaulay, and T. Zeuthen. 2009. ‘Ammonium ion transport by the AMT/Rh homolog TaAMT1;1 is stimulated by acidic pH’, Pflugers Arch, 458: 733–43.

Soliman, Moomen, and Ahmed Eldyasti. 2018. ‘Ammonia-Oxidizing Bacteria (AOB): opportunities and applications—a review’, Reviews in Environmental Science and Bio/Technology, 17: 285–321.

Steenwyk, J. L., T. J. Buida, Y. N. Li, X. X. Shen, and A. Rokas. 2020. ‘ClipKIT: A multiple sequence alignment trimming software for accurate phylogenomic inference’, Plos Biology, 18.

Stein, L. Y., D. J. Arp, P. M. Berube, P. S. Chain, L. Hauser, M. S. Jetten, M. G. Klotz, F. W. Larimer, J. M. Norton, H. J. Op den Camp, M. Shin, and X. Wei. 2007. ‘Whole-genome analysis of the ammonia-oxidizing bacterium, Nitrosomonas eutropha C91: implications for niche adaptation’, Environ Microbiol, 9: 2993–3007.

Stein, L. Y., and M. G. Klotz. 2011. ‘Nitrifying and denitrifying pathways of methanotrophic bacteria’, Biochem Soc Trans, 39: 1826–31.

Strassert, J. F. H., I. Irisarri, T. A. Williams, and F. Burki. 2021. ‘Author Correction: A molecular timescale for eukaryote evolution with implications for the origin of red algal-derived plastids’, Nat Commun, 12: 3574.

Stueeken, Eva E. 2020. ‘Hydrothermal vents and organic ligands sustained the Precambrian copper budget’, Geochemical Perspectives Letters.

Stüeken, Eva E. 2013. ‘A test of the nitrogen-limitation hypothesis for retarded eukaryote radiation: Nitrogen isotopes across a Mesoproterozoic basinal profile’, Geochimica et Cosmochimica Acta, 120: 121–39.

Stüeken, Eva E, Michael A Kipp, Matthew C Koehler, and Roger Buick. 2016. ‘The evolution of Earth’s biogeochemical nitrogen cycle’, Earth-Science Reviews, 160: 220–39.

Tavormina, P. L., V. J. Orphan, M. G. Kalyuzhnaya, M. S. Jetten, and M. G. Klotz. 2011. ‘A novel family of functional operons encoding methane/ammonia monooxygenase-related proteins in gammaproteobacterial methanotrophs’, Environ Microbiol Rep, 3: 91–100.

Thomazo, Christophe, Estelle Couradeau, and Ferran Garcia-Pichel. 2018. ‘Possible nitrogen fertilization of the early Earth Ocean by microbial continental ecosystems’, Nat Commun, 9: 2530.

Ullmann, R. T., S. L. Andrade, and G. M. Ullmann. 2012. ‘Thermodynamics of transport through the ammonium transporter Amt-1 investigated with free energy calculations’, J Phys Chem B, 116: 9690–703.

van Kessel, M. A., D. R. Speth, M. Albertsen, P. H. Nielsen, H. J. Op den Camp, B. Kartal, M. S. Jetten, and S. Lucker. 2015. ‘Complete nitrification by a single microorganism’, Nature, 528: 555–9.

Wagner, Florian B, Peter Borch Nielsen, Rasmus Boe-Hansen, and Hans-Jørgen Albrechtsen. 2016. ‘Copper deficiency can limit nitrification in biological rapid sand filters for drinking water production’, Water Res, 95: 280–88.

Wang, S., and H. Luo. 2021. ‘Dating Alphaproteobacteria evolution with eukaryotic fossils’, Nat Commun, 12: 3324.

Wang, S., and H. Luo. 2023. ‘Dating the bacterial tree of life based on ancient symbiosis’, bioRxiv: 2023.06.18.545440.

Ward, L. M., D. T. Johnston, and P. M. Shih. 2021. ‘Phanerozoic radiation of ammonia oxidizing bacteria’, Scientific Reports, 11.

Weidinger, K., B. Neuhauser, S. Gilch, U. Ludewig, O. Meyer, and I. Schmidt. 2007. ‘Functional and physiological evidence for a Rhesus-type ammonia transporter in Nitrosomonas europaea’, Fems Microbiology Letters, 273: 260–67.

Wijma, H. J., G. W. Canters, S. de Vries, and M. P. Verbeet. 2004. ‘Bidirectional catalysis by copper-containing nitrite reductase’, Biochemistry, 43: 10467–74.

Yang, Y., C. Zhang, T. M. Lenton, X. Yan, M. Zhu, M. Zhou, J. Tao, T. J. Phelps, and Z. Cao. 2021. ‘The Evolution Pathway of Ammonia-Oxidizing Archaea Shaped by Major Geological Events’, Mol Biol Evol, 38: 3637–48.

Yang, Z. H. 2007. ‘PAML 4: Phylogenetic analysis by maximum likelihood’, Molecular Biology and Evolution, 24: 1586–91.

Yool, A., A. P. Martin, C. Fernandez, and D. R. Clark. 2007. ‘The significance of nitrification for oceanic new production’, Nature, 447: 999–1002.

Zerkle, A. L., S. W. Poulton, R. J. Newton, C. Mettam, M. W. Claire, A. Bekker, and C. K. Junium. 2017. ‘Onset of the aerobic nitrogen cycle during the Great Oxidation Event’, Nature, 542: 465–67.

Zhang, H., S. Wang, T. Liao, Sean A. Crowe, and H. Luo. 2023. ‘Emergence of Prochlorococcus in the Tonian oceans and the initiation of Neoproterozoic oxygenation’, bioRxiv: 2023.09.06.556545.

Zorz, J. K., J. A. Kozlowski, L. Y. Stein, M. Strous, and M. Kleiner. 2018. ‘Comparative Proteomics of Three Species of Ammonia-Oxidizing Bacteria’, Front Microbiol, 9: 938.

Zuckerkandl, E, L Pauling, M Kasha, and B Pullman. 1962. ‘Horizons in biochemistry’, Horizons in Biochemistry: 97–166.

