## Supplementary Text and figures for "Dating ammonia-oxidizing bacteria with abundant eukaryotic fossils"

**This PDF file includes:**

Supplementary Text

Figures S1 to S6

References

#### Supplementary Text


**1 Data retrieval**

*1.1 xmoA homologs*

*1.2 Genome sequences*

**2 Phylogenetic analysis**

*2.1 xmoA gene tree based on amplicons and genomic sequences*

*2.2 Concatenated xmoCAB gene tree based on genomic sequences*

*2.3 Phylogenomic tree based on xmoA-containing genomes*

*2.4 Taxon sampling and the phylogenomic tree for molecular dating analysis*

**3 Molecular dating analysis**

*3.1 Four independent gene sets*

*3.2 Topological constraints for molecular dating analysis*

*3.3 Calibrations for molecular clocks*

*3.4 The impact of genes with large phylogenetic inconsistency*

*3.5 Bayesian sequential dating analysis*

**4 Comparative genomics analysis**

*4.1 Genome sampling and annotations*

*4.2 Identification of amoCAB-associated genes*

*4.3 Other amoCAB-associated genes*

#### 1. Data retrieval

##### *1.1 xmoA homologs*

We obtained 85,949 protein sequences of *xmoA*-like sequences from the NCBI *nr* database (accessed on 13 July 2021) by BLAST (2.9.0) against 42 reference sequences of *xmoA* (Dataset S1.2) with an E-value cutoff of 1e-20 and a coverage ratio over 60%. To ensure the quality of the obtained *xmoA* homologs, we filtered out 3,101 sequences that were not annotated as *xmoA* (K10944) using KofamScan (updated in 2019) which comprises 25,614 predefined probabilistic models each for a distinct functional gene family (Aramaki, et al. 2020), based on the adaptive thresholds for each model defined by KEGG. The interconnectivity of NCBI's taxonomy, genome, and protein databases was identified via the Entrez search (Kans 2020) and custom Python scripts (see Data and Code availability), which enable us to collect the taxonomic information and the source genome, if any, of the collected *xmoA* homologs. The collected *xmoA* homologs were mainly from the phyla Proteobacteria, Nitrospirota, Actinobacteria, Verrucomicrobia, the candidate division NC10, and Thaumarchaeota. Complete information on the collected *xmoA* homologs is available in the online repository (Dataset S1.4).

The collection of reference *xmoA* is used to comprehensively capture the phylogenetic diversity of the gene that encodes copper-containing membrane monooxygenases (CuMMO). A total of 42 *xmoA* sequences (Dataset S1.2), which encode CuMMO with varying substrate preference, were obtained from several studies (Tavormina, et al. 2011; Coleman, et al. 2012; Kozlowski, et al. 2016; Boddicker and Mosier 2018; Khadka, et al. 2018). The reference gene set includes five types of *xmo*: *pmo* encoding particulate methane monooxygenase, *bmo* encoding butane monooxygenase, *emo* encoding ethene monooxygenase, *pxm* encoding alternate methane monooxygenase, and *amo* encoding ammonia monooxygenase. Note that our study did not include the soluble methane monooxygenase, which belongs to a different protein family rather than CuMMO (Banerjee, et al. 2015).

##### *1.2 Genome sequences*

A large portion of the 82,848 *xmoA* homologous sequences were derived from amplicon sequencing. Only 1,111 *xmoA* homologs were associated with 880 genome sequences (Dataset S1.1). We expanded our genome set by including 1) the ammonia-oxidizing bacteria (AOB) or complete ammonia oxidizing bacteria (CAOB) that were not included in the above and 2) the phylogenetically related lineages that are not AOB or CAOB. For Nitrospirota where CAOB belong to (Daims, et al. 2015), we downloaded all 697 Nitrospirota genomes from GenBank (accessed on 13 July 2021) (Benson, et al. 2013). In terms of Proteobacteria where AOB belong to, downloading all 604,712 genomes incurred a huge computational burden. We noted that in Proteobacteria a few genera each contain numerous genomes (e.g., over 50,000 genomes in *Salmonella* and *Escherichia*), whereas over 90% of the genera contain fewer than 30 genomes. We therefore downloaded two "representative genomes," which are defined by the NCBI, from each of the genera that contain over 30 genomes but all the genomes of the remaining genera. The exceptions are the families Nitrococcaceae (Gammaproteobacteria) and Nitrosomonadaceae

(Betaproteobacteria) where the proteobacterial AOB belong to (Lehtovirta-Morley 2018; Stein 2019). We downloaded all the genomes from these families to mitigate any bias in taxon selection. The related scripts are available in the online repository (see Data and Code availability). This procedure retrieved 6,585 (Dataset S1.1).

Apart from the bacterial genomes that contain *xmo* and their phylogenetic relatives, we downloaded eukaryotic genomes for molecular dating analysis with the aid of eukaryotic fossils. We followed Wang and Luo (2021) and Zhang, et al. (2023) to obtain 34 plastid, 16 mitochondrial, and 25 eukaryotic nuclear genomes (Dataset S1.7). We also downloaded two additional cyanobacterial genomes from the genus *Gloeomargarita* which is the closest cyanobacterial lineage to the eukaryotic plastid lineage (Ponce-Toledo, et al. 2017). Likewise, we downloaded 10 additional alphaproteobacterial genomes covering seven orders that are related to the mitochondrial lineage and were used in the previous study (Wang and Luo 2021; Liao, et al. 2022). Including these lineages helps constrain the ages of eukaryotic lineages based on their plastid and mitochondrial genomes. Finally, we downloaded 21 high-quality genomes of oxygenic Cyanobacteria and two genomes of Vampirovibrionia (formerly known as Melainabacteria) from RefSeq and GenBank (Benson, et al. 2013) for the implementation of bacterial calibration, as done in the previous studies (Matheus Carnevali, et al. 2019; Liao, et al. 2022).

In total, the compiled genome set included 7,627 bacterial genomes (Dataset S1.1) and 75 eukaryotic genomes (Dataset S1.7). All bacterial genomes without annotations were subjected to annotation using Prokka (v1.14.5) (Seemann 2014). The estimation of genome completeness was conducted using miComplete (v1.1.1) (Hugoson, et al. 2020). The protein sequences of eukaryotic genomes were obtained from their source databases (Dataset S1.7).

#### 2. Phylogenetic analysis

We employed IQ-Tree (v1.6.12) (Nguyen, et al. 2015) to determine the best-fitting amino acid substitution model (for both gene tree and genome tree construction) and data partition model (for genome tree construction), and to generate the phylogeny based on that model. The protein sequence-based phylogenies were constructed with the parameters '-mset WAG,LG,JTT,Dayhoff -mrate E,I,G,I+G -mfreq FU' and 1,000 replicates of the ultrafast bootstrap. The generated phylogenies were visualized using iTOL v5 (Letunic and Bork 2021).

##### *2.1 xmoA gene tree based on amplicon and genomic sequences*

Using CD-HIT (v4.8.1) (Fu, et al. 2012) with the default sequence similarity cutoff (0.9), we grouped the 82,848 putative *xmoA* homologs into 881 clusters. Together with 42 curated *xmoA* as references (Dataset S1.2), a total of 923 *xmoA* were aligned at the amino acid level using the G-INS-I refinement method in MAFFT (v7.471) (Katoh and Standley 2013). Uninformative sites in the alignment were removed using ClipKit (v1.1.5) with the smart gap mode, which automatically determines the gap threshold for trimming (Steenwyk, et al. 2020). The gene tree was subsequently constructed by IQ-Tree. The best-fit model WAG+G4 was selected based on the Bayesian Information Criterion (BIC) score. The generated tree was rooted at the archaeal *amoA* lineage following the previous study (Khadka, et al. 2018). Apart from the outgroup-based rooting method, an outgroup-free method, namely minimum variance (MV v1.5) (Mai, et al. 2017) was employed. This method uses branch lengths as an input to infer the optimal root position by minimizing the cost based on its deviation from the strict molecular clock. Both the outgroup-based method and the MV method inferred the same branch, leading to the ancestor of actinobacterial *xmoCAB* and proteobacterial *xmoCAB*. The earliest branching lineage in the gene tree comprising *amoCAB* genes from Beta-AOB and CAOB is different from that in the concatenated *xmoCAB* gene tree (Fig. 1A). This difference is likely caused by the insufficient phylogenetic information in single genes (Philippe, et al. 2011).

##### *2.2 Concatenated xmoCAB gene tree based on genomic sequences*

We also constructed the gene phylogeny based on the concatenated *xmoCAB* derived from 848 *xmoA*-containing genomes (Dataset S1.3). The remaining 11 genomes were not included because they contained *xmoA* but lacked *xmoBC*. Each subunit gene was aligned by MAFFT (v7.471) using the G-INS-I refinement method (Katoh and Standley 2013). All the three alignments were then concatenated for phylogenetic tree reconstruction (Fig. 1A) using IQ-Tree (v1.6.12) with the option '-spp' enabling partitioned analysis, which allows each partition to have independent rates. The archaeal *xmoCAB* sequences which showed large phylogenetic distances to the bacterial *xmoCAB*, were removed to avoid long-branch attraction artifacts (Bergsten 2005). Following that, the generated tree was rooted using the *bmo* lineage as an outgroup, which was used in a previous study (Khadka, et al. 2018). The outgroup-free method, MV, placed the root at the same phylogenetic position as the outgroup-based method (Mai, et al. 2017).

*2.3 Phylogenomic tree based on xmoA-containing genomes*

The phylogenomic tree (Fig. 1B) of the *xmoA*-containing genomes was generated using PhyloPhlAn (v3.0) (Segata, et al. 2013), which is an integrated pipeline that includes phylogenomic marker gene identification, sequence alignment, alignment trimming and tree construction. The diversity parameter in the PhyloPhlAn was set as ‘high’, given that the *xmoA*-containing genomes were sourced from different phyla. This parameter allows for simultaneous configuration of parameters for several steps, such as trimming, subsampling and selection of marker gene sets. PhyloPhlAn subsequently excluded the genomes that contained fewer than 100 out of the 400 marker genes identified by this tool. As a result, 848 genomes remained. To increase the confidence that the retrieved *amoA*-containing genomes are capable of ammonia oxidation, we further annotated the genes, *cycAB*, which encode the necessary electron carriers for ammonia oxidation (Gonzalez-Cabaleiro, et al. 2019; Picone, et al. 2021) by BLASTP with an E-value threshold of 1e-20. The quinone-reducing tetraheme cytochrome $c_M552$ (*cycB*), soluble cytochrome *c554* (*cycA*), and the hydroxylamine oxidoreductase (*hao*) constitute the hydroxylamine-ubiquinone redox module (HURM), which is a unique cyclic electron flow module found in the AOB (Arp, et al. 2007; Klotz and Stein 2008). The gene *hao* was not included in this study since the complex evolution history and extensive duplications (Klotz, et al. 2008) may confound AOB identifications.

*2.4 Taxon sampling and the phylogenomic tree for molecular dating analysis*

As mentioned above, we compiled 7,683 genomes including *xmoA*-containing genomes and their relatives (Dataset S1.1; Section 1.2). The predicted protein sequences by Prokka (v1.14.5) were subjected to annotation using HMMER (v3.2.1) (Mistry, et al. 2013) based on the pre-built HMM profiles of the 120 bacterial marker proteins (bac120) (Dataset S1.5) that were suggested to be useful for the bacterial tree inference (Parks, et al. 2017), with an E-value cutoff of 1e-50. The genomes with fewer than 24 (20%) out of the 120 genes were removed. This resulted in a final dataset containing 7,037 genomes.

The taxon sampling method built upon the "prototype selection" strategy was developed in the previous study (Zhu, et al. 2019), which takes the final number of genomes to be retained and pairwise distance calculated by Mash (Ondov, et al. 2016) as inputs. We implemented a strategy that considered both maximizing the phylogenetic distance between genomes and preserving basal phylogenetic lineages without the need to predetermine the final number of genomes. First, we used Mash (v2.2), which is a k-mer based metric, to compute pairwise similarities between genomes using a sketch size of 10,000 k-mers, with each consisting of 22 bp based on the default setting. Then, we used the pairwise similarities to perform the selection. The strategy involved the selection of two sets of genomes: 1) representative genomes from the focal lineage, and 2) the genomes that help to determine the total group of the focal lineage and maximize the phylogenetic diversity.

The first set of genomes was determined based on the pairwise distance from mash, followed by unsupervised clustering using density-based spatial clustering of applications with noise (DBSCAN) (Ester, et al. 1996; Schubert, et al. 2017). The three ammonia-oxidizing lineages, Beta-AOB, Gamma-AOB, and CAOB (clades represented in red branches in Fig. 1B), which were determined based on the presence of *cycAB* (Fig. 1B) and the phylogenetic positions in the *xmoCAB* phylogeny (Fig. 1A), were chosen as the focal lineages. In addition, with respect to the significant proportion of genomes in Gammaproteobacteria containing *pmoCAB* (Fig. 1B), which is indicative of the importance of methane-oxidizing bacteria (MOB), the Gamma-MOB lineage was also chosen as a focal lineage. Therefore, the implementation of Mash on these focal lineages resulted in 61, 5, 39 and 103 clusters for Beta-AOB, Gamma-AOB, CAOB, and Gamma-MOB, respectively. For each cluster, one representative genome with the highest genome completeness was chosen. To avoid selecting too many genomes and thus increasing the computational cost of the subsequent molecular dating analysis, five early branching and representative genomes were manually selected for the Beta-AOB, CAOB, and Gamma-MOB, respectively.

The second set of genomes was chosen based on the mash distance between the genomes of the focal lineages and the remaining genomes from the abovementioned 7,683 genomes. We sampled genomes from the remaining genome pool for each focal lineage using two specific criteria. The first criterion was based on the smallest mash distances, while the second criterion involved selecting genomes that were not distributed within the focal lineage in the phylogenomic tree (Fig. 1B). Initially, we sampled 500 non-focal genomes for each of the 5, 61, 39, and 103 representative genomes of Gamma-AOB, Beta-AOB, CAOB, and Gamma-MOB, respectively. Thus, we would in theory sample 2,500, 30,500, 19,500, and 51,500 non-focal genomes, respectively. However, the actual number of non-focal genomes we sampled was 336, 300, 344, and 361, respectively, because of the repeated sampling of some of the non-focal genomes. Since these repeatedly sampled non-focal genomes usually showed higher similarity and thus closer evolutionary relationship to the focal genomes, we further shortlisted the non-focal genomes to the ten most frequently sampled. Collectively, we selected a total of 63 genomes from the Proteobacteria phylum where Gamma-AOB, Beta-AOB, and Gamma-MOB belong to, and the Nitrospirota phylum where CAOB belong to.

As we aim to compare the origin time of AOB and anaerobic ammonia oxidation bacteria (AnAOB) that perform aerobic and anaerobic ammonia oxidation, respectively, we also sampled five genomes of AnAOB, four genomes of evolutionarily related bacteria in Planctomycetes where AnAOB belong to, and one genome in Verrucomicrobiota according to our recent study (Liao, et al. 2022). Together with the ten Alphaproteobacteria, 21 oxygenic Cyanobacteria, and two Vampirovibrionia (formerly known as Melainabacteria) that serve as the most closely related lineages to oxygenic Cyanobacteria (Section 1.2), a total of 106 bacterial genomes were sampled for the following molecular dating analysis.

#### 3. Molecular dating analysis

We compiled four independent gene sets (Dataset S1.5) for molecular clock analysis. The traditional approach to performing molecular dating analysis involves the utilization of bacterial fossils and calibrations mainly in Cyanobacteria and requires the identification of gene sets that are conserved across focal bacterial lineages and the Cyanobacteria. Recently, alternative sets of bacterial genes that are conserved in eukaryotic genomes based on mitochondrial (Munoz-Gomez, et al. 2019; Munoz-Gomez, et al. 2022) and plastid endosymbiosis (Ponce-Toledo, et al. 2017) have been identified, which allows using the rich eukaryotic fossils to calibrate bacterial evolution (Wang and Luo 2021; Zhang, et al. 2023). In the present study, we sampled nuclear, mitochondrial, and plastid genomes of eukaryotes based on the abovementioned studies, and implemented the molecular dating analysis using the program MCMCTree from the PAML package (4.9j) (Yang 2007). In the following sections, we detailed the gene sets and the fossil calibrations in molecular clock analysis. In all clock analyses, two replicates were conducted for each run with identical parameters (burn-in: 60,000; sample frequency: 30; number of samples: 20,000) to ensure convergence (Yang and Rannala 2006; Rannala and Yang 2007).

##### *3.1 Four independent gene sets*

Four independent gene sets (Dataset S1.5) were used in the molecular clock analysis. The first gene set, referred to as Battistuzzi25, contains the 25 genes universally conserved among bacterial genomes (Battistuzzi and Hedges 2009). The molecular clock analysis based on this gene set did not involve eukaryotic genomes. The second gene set, namely ‘Mito24’, consists of the 24 genes conserved between alphaproteobacterial and mitochondrial genomes (Wang and Wu 2015). Here, 16 eukaryotic mitochondrial genomes were included. The third gene set, ‘Gomez19’, consists of 19 mitochondrion-originated genes that were likely transferred to the eukaryotic nuclear genome and that are conserved across most bacterial lineages. Briefly, Munoz-Gomez, et al. (2022) identified 108 eukaryotic genes with an alphaproteobacterial origin, from which Wang and Luo (2023) identified 19 genes that were conserved across the bacterial tree of life, thus a gene set suitable for molecular clock analysis with the strategy using eukaryotes’ fossils to date the evolution of distantly related bacterial lineages. Accordingly, 25 nuclear genomes of eukaryotic organisms were chosen, following a recent study (Wang and Luo 2021). The fourth gene set ‘Plastid39’ includes 39 genes that are conserved between plastid and cyanobacterial genomes (Ponce-Toledo, et al. 2017). The molecular dating analysis based on it incorporated 36 plastid-containing eukaryotes. The eukaryotic genomes used in the molecular dating analysis were provided in Dataset S1.7.

To find the orthologous genes of the four gene sets, we conducted HMMER v3.2.1 (Mistry, et al. 2013) or BLASTP 2.9.0 (Johnson, et al. 2008) analysis using an E-value threshold of 1e-20 against the 106 bacterial genomes (see Section 2.4). For the BLASTP search, the coverage ratio of the subject protein was required to be over 60%. The selection of the two methods, BLASTP or HMMER, was based on the type of reference gene. For the reference sequences sourced from Pfam (El-Gebali, et al. 2019), HMMER was used for protein annotation. For the remaining reference sequences, BLASTP was used. Only those genes that were identified in more than 20% of the 106 bacterial genomes were used in molecular dating analyses (Dataset S1.8).

*3.2 Topological constraints for molecular dating analysis*

The molecular dating analysis implemented in MCMCTree requires a fixed topology as an input. To enable the molecular clock analysis with the gene set “Battistuzzi25”, in which eukaryotes were not included, the phylogenomic tree topology of the 106 bacterial genomes (Section 2.4) was used. We constructed a maximum likelihood phylogenomic tree based on the concatenated alignment of the retrieved “bac120”proteins (Dataset S1.5), with the parameters ‘-mset WAG, LG, JTT, Dayhoff -mrate E,I,G,I+G -mfreq FU -m TESTONLY -madd LG+C20+G, LG+C30+G, LG+C40+G, LG+C50+G, LG+C60+G’ and 1,000 replicates of the ultrafast bootstrap (Minh, et al. 2013). Here, the parameters ‘-madd LG+C20+G, LG+C30+G, LG+C40+G, LG+C50+G, LG+C60+G’ instructed the IQ-Tree to search across five complex amino acid substitution models that have 20, 30, 40, 50, and 60 categories of amino acid frequency profiles, respectively (Quang le, et al. 2008). These profiles were learned from a large and curated alignment databases (Sander and Schneider 1991) and benchmarked against another independent database (Sanderson, et al. 1993). The model LG+C60+G was selected as the best-fit model based on the model test (lowest AIC by default) by ModelFinder (Kalyaanamoorthy, et al. 2017) and employed with the posterior mean site frequency (PMSF) approximation (Wang, et al. 2018), which accounts for the site-specific amino acid equilibrium frequencies thus amino acid preferences for each site, as implemented in IQ-Tree (v1.6.12).

Molecular clock analyses with the remaining three gene sets each involved eukaryotes and thus required a different topological structure. All these topologies built on the topology of the 106 bacterial genomes but also included eukaryotic genomes and additional bacterial genomes. In sum, the trees used as topological constraints for “Plastid39”, “Gomez19”, and “Mito24” contained 142, 131, and 122 genomes, respectively (Fig. 1D-1G). Each of these three topological structures had a bacterial portion and a eukaryotic portion. The topological structures of the bacterial portion were generated by using the abovementioned 106-genome tree as the guide tree and taking the alignment of bac120 proteins across the expanded bacterial genome lists. The topological structures of eukaryotic lineages were obtained from previous studies (Wang and Luo 2021; Zhang, et al. 2023) and manually grafted to those of the corresponding bacterial lineages. These species tree topology was available in the online repository.

##### *3.3 Calibrations for molecular clocks*

For the “Battistuzzi25” strategy which does not involve eukaryotic lineages, only bacterial calibrations can be used. We included four bacterial calibrations: one geochemical constraint related to GOE that was used to calibrate the total group of the oxygen-evolving Cyanobacteria, two molecular fossils targeting two cyanobacterial lineages, and one molecular biomarker targeting purple sulfur bacteria. In addition, a root maximum age is required (Yang and Rannala 2006). These are detailed as follows.

For the root of the tree, there is no solid evidence to constrain the timing of the earliest bacteria. We therefore implemented several possible maximum constraints from 3.5

Ga to 4.5 Ga. The 3.5 Ga is based on the stromatolites of the Pilbara Supergroup (3,490 - 3,240 Ma), which provides the oldest evidence of life on Earth (Van Kranendonk 2006). The 4.5 Ga is based on the age of the Earth (Barry and Taylor 2013). For the total group of oxygenic Cyanobacteria (BN2 in Fig. 1G), we set 2,300 Ma as the minimum boundary (Bekker, et al. 2004). Since the maximum boundary of the time constraints is hard to determine (Zhang, et al. 2021), we did not impose any maximum boundary for the total group of oxygenic Cyanobacteria (calibration sets B1 to B5). For the remaining two calibration nodes in Cyanobacteria, we followed Zhang, et al. (2021) and set 1,700 Ma as the minimum boundary for the total group of Pleurocapsales (BN4 in Fig. 1G) based on the presence of microfossils (Golubic and Lee 1999) and set 1,600 Ma as the minimum boundary for the total group of Nostocales (BN3 in Fig. 1G) based on the discovery of the Nostocalean Akinetes fossil (Golubic, et al. 1995). Two alternative ages of Nostocales, 1,200 Ma (Horodyski and Donaldson 1980) (calibration sets B7 and B8) and 2,000 Ma (Amard and Bertrand-Sarfati 1997) (calibration set B5), were also set as the minimum boundary for comparisons. Moreover, we set 1,640 Ma as the minimum boundary for the total group of Chromatiaceae (a family in Gammaproteobacteria where one AOB lineage is affiliated; BN5 in Fig. 1G) based on the biomarker evidence sourced from the purple sulfur bacteria found in the 1.6 Ga basin (Brocks, et al. 2005).

For the "Mito24" strategy utilizing eukaryotic fossils, we imposed four eukaryotic fossil-based calibrations. These calibrations (Fig. 1D) were implemented following the Supplementary Note S3.2.1 in Wang and Luo (2023), including the crown group of

Angiosperm (XN1), the crown group of Embryophyta (XN2), the total group of Florideophyceae (XN3), and the crown group of Rhodophyta (XN4). We did not use the alternative calibration scheme of the node XN4, such as 1.2 Ga, in our study, as these alternative calibration schemes had small influence on posterior age estimates (Wang and Luo 2023).

For the "Gomez19" strategy, we incorporated a total of nine eukaryotic fossil-based time constraints (Fig. 1E) derived from Supplementary Note S3.2.1 in Wang and Luo (2023). In addition to the same calibrations utilized in the Mito24 gene set (XN1 to XN4), we used five more calibrations, including the crown group of Amniota (AN1), the crown group of Chordata (AN2), the crown group of Metazoa (AN3), the total group of Fungi (AN4), and the crown group of Dikarya (AN5). The time constraints of the above nine nodes were set by following Wang and Luo (2023).

For the "Plastid39" strategy, we incorporated a total of nine eukaryotic fossil-based time constraints (Fig. 1F) derived from our recent study (Zhang, et al. 2023). In addition to the same fossils and calibrations utilized in the Mito24 gene set (XN1 to XN4), we used five other fossils including the total group of Charophyta (PN1), the crown group of Chlorophyta (PN2), the crown group of Spermatophyta (PN3), the crown group of Pelagophyceae (PN4), and the crown group of Diatom (PN5).

It should be noted that the calibrations implemented based on the four bacterial calibrations (Fig. 1G) mentioned above were utilized in combination with the eukaryotic fossil-based calibration for the strategies "Gomez19", "Mito24", and "Plastid39". For example, the calibration set denoted as B1P suggests the amalgamation of the set B1 (see Dataset S2.1) of bacterial fossil-based calibration (BN1: 4.5-2.32 Ga, BN2: > 2.32 Ga, BN3: 4.5-1.6 Ga, BN4: 4.5-1.7 Ga, BN5: 4.5-1.6 Ga) and the eukaryotic fossil-based calibration set P (see Dataset S2.1) (XN1: 2.50-1.25 Ga, XN2: 5.09-4.55 Ga, XN3: 18.91-5.25Ga, XN4: 18.91-10.45 Ga, PN1: 18.91-4.8 Ga, PN2: 18.91-10.56 Ga, PN3: 5.09-3.08 Ga, PN4: 18.91-1.34 Ga, PN5: 18.91-0.08 Ga) for "Plastid39".

##### *3.4 The impact of genes with large phylogenetic inconsistency*

The genes used in molecular dating analyses may have different evolutionary histories from the organisms, which can affect the posterior age estimates. Here, we evaluated the incongruence between the gene tree and the species tree using the delta log-likelihood (ΔLL), a metric used in several phylogenetic studies (Shen, et al. 2017; Smith, et al. 2020). On this basis, two different strategies were developed to evaluate the effect of removing the most incongruent genes on the posterior time estimates.

To implement the two strategies, ΔLL was computed by comparing the likelihood values of the gene tree that reconstructed with and without the reference tree topology using IQ-Tree (v1.6.12) (Shimodaira 2002). The gene tree was constructed based on the amino acid alignment using IQ-tree with the parameters '-m LG -redo -wbtl -bb 1000'. These two trees were taken as input trees to compute the phylogenetic incongruence, delta log likelihood (ΔLL), by the IQ-Tree with the approximately unbiased (AU) test (Shimodaira 2002) and RELL approximation (Kishino, et al. 1990).

One strategy, namely the "gene-removal" approach, gradually removing the most incongruent genes with the largest ΔLL and generating a series of subsets of molecular data (Fig. S1). The differences in time estimates by using these datasets may result from two processes: the decrease of phylogenetic information due to the use of fewer genes and the increase of phylogenetic congruence among the retained genes due to the use of the remaining genes showing reduced ΔLL (Fig. 2B). To disentangle them, we developed a "sliding window-based" gene selection strategy (Fig. 2C), where we first sorted out the genes according to ΔLL and then iteratively selected a subset of ten genes with decreasing ΔLL. By performing the molecular clock analysis with a fixed number of genes, the differences in age estimates are largely attributed to the differences in ΔLL between the selected subsets of genes. The use of the datasets derived from gene-removal approach estimated comparable origin time of ammonia oxidizers (Fig. 2A). While the gene-removal approach removes genes with high phylogenetic incongruence, it also decreases the amount of phylogenetic information available for dating analysis. Both factors are likely responsible for the divergent age estimations of the four focal lineages (Fig. 2B).

Both the "sliding window" and "gene removal" strategies require many separate dating analyses, thereby incurring a high computational expense. Using fewer partitions in the dating analysis, which was accomplished by clustering genes and concatenating the genes from the same cluster into a single partition, is able to alleviate this issue. This approach is supported by the observation that age estimates derived from the timing analysis using partitions greater than three are similar to those derived from that using more partitions

(Fig. S2). The linear correlation analysis conducted between age estimates using the full partitions (39 partitions for Plastid39) and those using the fewer partitions (ranging from 1 to 36) yielded coefficients of determination ($R^2$) ranging from 0.95 to 0.9998. As $R^2$ increases, age estimates using fewer partitions become increasingly comparable to those using the full partitions (Fig. S2). In the present study, three partitions, whose $R^2$ reached 0.9755, were used for each MCMCTree analysis. To partition the genes, we followed Angelis, et al. (2018) by clustering the genes based on the average evolutionary rates of each gene, which is computed by codeml implemented in PAML (Yang 2007). The K-means clustering algorithm was subsequently employed since it can generate a given number of clusters.

##### *3.5 Bayesian sequential dating analysis*

The Bayesian sequential strategy (Dos Reis, et al. 2012) has recently been employed to date the mammalian evolutionary tree (Álvarez-Carretero, et al. 2022). It is a two-step procedure. The first step performs a clock analysis on a species tree topology with fewer taxa, followed by a second step that performs another clock analysis on an expanded species tree with more taxa and uses the posterior age estimates derived from the first step as the priors. Using the posterior age estimates from the first step as the time priors in the second step is able to better constrain the expanded phylogenomic tree. Note that the dating software used here, MCMCTree, acknowledges five different types of distributions as time constraints, namely the gamma, skew-normal, skew-t, Cauchy, and uniform distributions. Among them, Cauchy and uniform distributions are generally not suitable to fit the posterior dates (they are usually used to fit prior calibrations which are generally less informative) (Wang and Luo 2023). Thus, for each node in the first-step species tree, the posterior ages were fitted with the three distributions using the function ‘fitdist’ implemented in the R package (fitdistplus v1.1-8) (Delignette-Muller and Dutang 2015). The log-likelihood value of the fit was used to calculate the Akaike Information Criterion (AIC) score and subsequently used to select the best fit distribution from the abovementioned three distributions (gamma, skew-normal, skew-t). Finally, the parameters of each fitted distribution are determined (Dataset S2.1) and used to set time constraints in the second-step dating analysis.

Importantly, the gene sets used in the two steps must be independent from each other (i.e., the same gene cannot be used twice) (Álvarez-Carretero, et al. 2022). Furthermore, the posterior ages from the first-step clock analysis that were used as time constraints in the second-step analysis must have non-overlapped distributions. Otherwise, the basic criterion that parent nodes must be older than their child nodes will truncate the imposed time constraints, and it would result in a poor approximation in the second-step analysis. For the first-step dating analysis (Fig. 3A), we included 40 nuclear genomes of eukaryotes (Dataset S1.7) and 320 orthologues that were previously selected for the purpose of dating the eukaryotic evolution (Strassert, et al. 2021). The 40 nuclear genomes analyzed here were the same eukaryotic lineages mentioned earlier (Section 3.3), with the exception of diatom nuclear genomes. As such, a total of 12 out of 14 calibrations nodes (XN1 to XN4, PN1 to PN3, and AN1 to AN5) were imposed, with the exclusion of two calibration nodes (PN4, PN5) (Fig. 3; calibration set EUK in Dataset S2.1). Moreover, we used nine partitions to reduce computational costs (Section 3.4). As shown in Figure 3A, 12 calibration nodes in the eukaryote tree did not overlap in the posterior ages with their parent and child nodes and thus were used as time constraints in the second-step clock analysis. We implemented sequential clock analyses with each of the three gene sets (Plastid39, Gomez19, and Mito24) that involve eukaryotic lineages. As introduced above (Section 3.4), we selected the three gene sets that removed 24, 6, and 18 genes, respectively, with the smallest ΔLL and used three partitions for the second-step clock analysis (Fig. S3). The age estimates of the five focal lineages (Gamma-AOB, Beta-AOB, CAOB, AnAOB, and eukaryotes) were shown in density plots (Fig. 3B).

#### 4. Comparative genomics analysis

##### *4.1 Genome sampling and annotations*

The genomes used for comparative genomic analysis was part of the genomes sampled for molecular clock analysis (see Section 2.4). We kept all genomes of the four focal lineages (Beta-AOB, Gamma-AOB, CAOB, and MOB) and their sister lineages. Together with all the genomes belonging to the focal lineages, a total of 1,119 genomes were included. The predicted proteins from these genomes were further annotated by HMMER against the KEGG database (Kanehisa and Goto 2000) with an E-value cutoff of 1e-20. Multidimensional scaling (MDS) was then performed based on the pairwise Manhattan distance matrix and the pairwise mash distance matrix (Figs. 4A & 4B), which represent metabolic similarity and nucleotide similarity, respectively. The two distance matrices for the 1,119 genomes are available in the online repository.

##### *4.2 Identification of amoCAB-associated genes*

To identify *amoCAB*-associated genes, we converted the presence and absence of genes (including *amoCAB* and KEGG-annotated genes) into binary values (1 and 0). Then we performed all three tests including Fisher-exact test, Pearson correlation test, and the phylogenetic signal test (phylosig) (Revell 2012) for each gene. Specifically, the fisher exact test examines the significance of the association between the distribution of *amoCAB* and tested genes. The Pearson test examines the linear correlation (either positive or negative) between the target genes. In addition to statistical tests based on numerical values, the phylosig method examines the signal (Blomberg, et al. 2003) of the genes along with the phylogenetic tree, which help to find the genes that emerged through convergent evolution or vertical inheritance.

For each gene, we performed Fisher-exact test and Pearson correlation between i) AOB, CAOB and others, ii) AOB and non-AOB proteobacteria, iii) Gamma-AOB and non-AOB gammaproteobacteria, iv) Beta-AOB and non-AOB betaproteobacteria, and v) CAOB and non-CAOB Nitrospirota. The phylosig method was conducted separately for each gene and the species tree. The obtained *p*-values were adjusted using the Benjamini-Hochberg false discovery rate (FDR) procedure. The genes with corrected p-values less than 0.05 in five groups of comparisons are manually examined according to the correlation coefficient and the phylogenetic signal derived from the R package “phylosig”. Only the genes with significant phylogenetic signal and large correlation coefficient are shown in the Figure 4. Other significant genes are available in the online repository.

*4.3 Other amoCAB-associated genes*

The mechanisms underlying the osmotic and pH adaptations in AOB have been previously documented (Lehtovirta-Morley 2018). For example, the gene (*torY*) encoding (Fig. 4) the reductase for trimethylamine N-oxide (TMAO) is used to regulate osmotic stress (Yin, et al. 2017). Furthermore, the gene *odc* encodes a protein in ornithine cycle, which help the organism to maintain a neutral cytoplasmic pH (Harden, et al. 2015). Furthermore, three genes implicated in stress tolerance have been identified. For instance, the *yqjG* gene, which encodes glutathione reductase, plays a pivotal role in detoxifying both internal and external toxic compounds, while also offering protection from oxidative stress (Green, et al. 2012) (Fig. 4). Similarly, the *ynfA* gene, encoding an SMR family efflux pump, which increases the antibiotics’ resistance of the host, is markedly prevalent in AOB and CAOB compared to their closest relatives (Sarkar, et al. 2015). The *spoIVFB* gene, likely assisting AOB and CAOB in forming spores, is involved in another key survival mechanism during severe environmental shifts (Cutting, et al. 1991). Additionally, genes that encode glycine dehydrogenase have been found to play a crucial role in maintaining the redox potential of the NAD pool under oxygen-limited conditions (Hutter and Dick 1998). In contrast to the presence of the gene *gldC* (glycine dehydrogenase) in both eukaryotes and prokaryotes (including CAOB), *gcvPA* and *gcvPB* (glycine dehydrogenase) are exclusively found in AOB. However, the roles of these two genes in AOB remains largely unresolved.

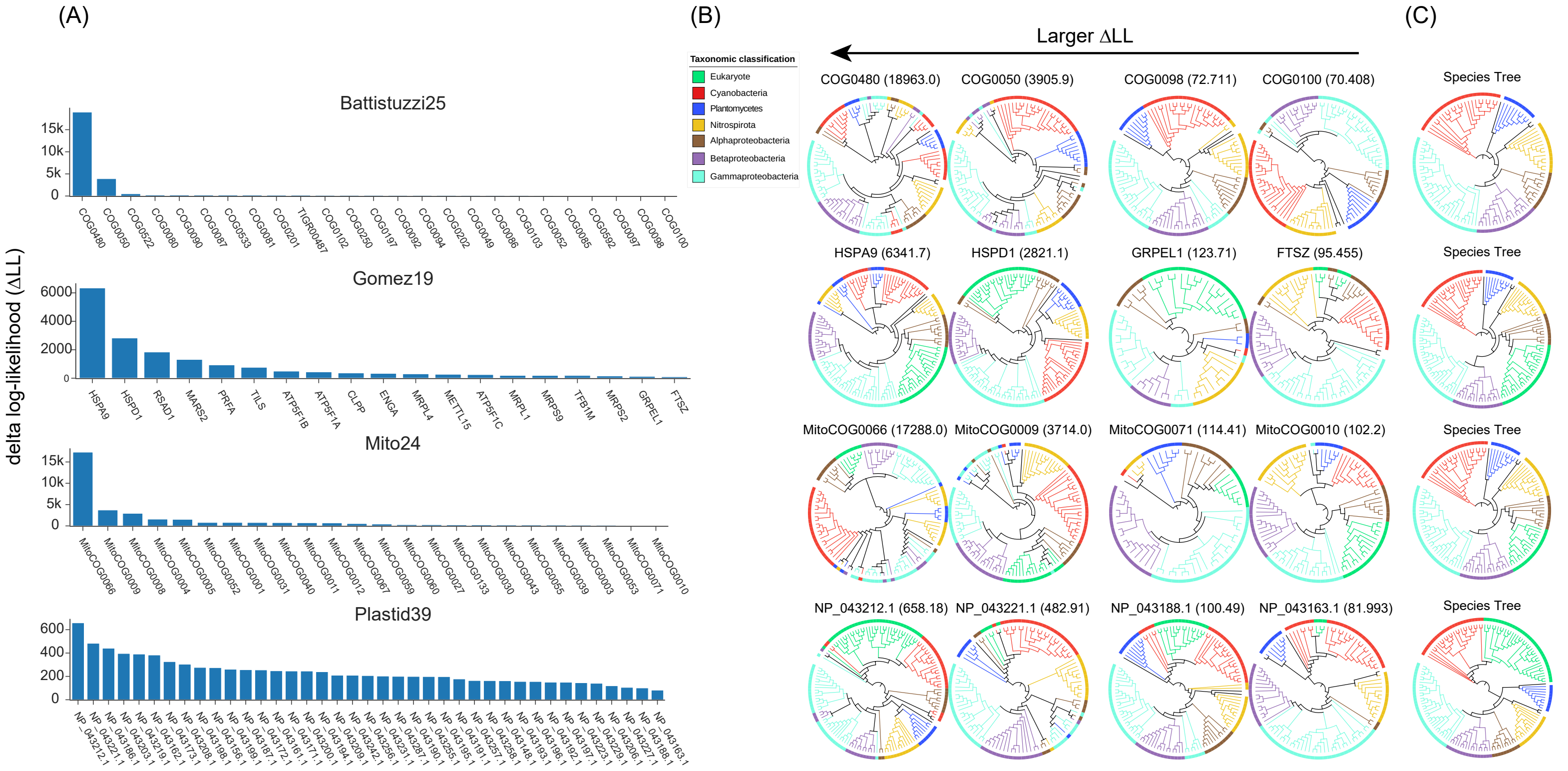


**Figure S1**. The delta log-likelihood (ΔLL) of genes in the four gene sets (Battistuzzi25, Gomez19, Mito24, and Plastid39). (A) The genes within each gene set are arranged in descending order based on their ΔLL. The larger the ΔLL value of a gene, the greater the incongruence between the gene tree and the species tree. (B) The four gene trees shown here for each gene set include the two genes with the largest ΔLL and the two genes with the smallest ΔLL. (C) The species trees. The colors of the branches and the strips next to the tips indicate the taxonomy information. The presence/absence patterns of each gene in different lineages are provided in the Dataset S1.8.

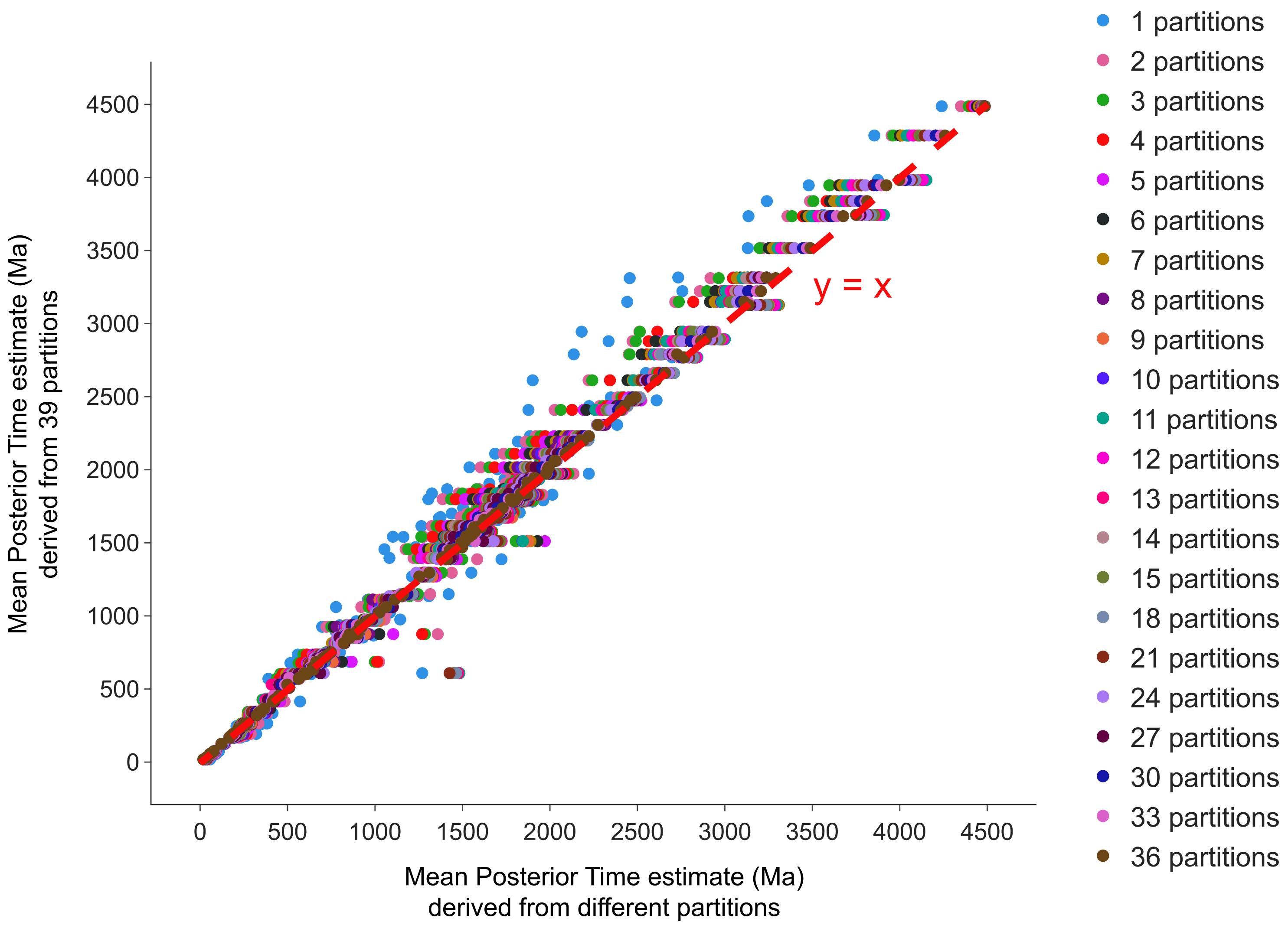


**Figure S2**. Scatter plot showing the differences in posterior age estimates between those derived from the 39 partitions (y-axis) and those derived from the fewer partitions (ranging from 1 to 36) (x-axis). Each circle represents an internal node of the species tree used in the dating analysis. The node colors correspond to the number of partitions used in the dating analysis. The nodes close to the red diagonal line represent the nearly identical age estimates that were derived from the analyses using fewer partitions and those using 39 partitions. The dating analysis shown here was performed with the Plastid39 gene set and the B1P calibration set (Dataset S2.1).

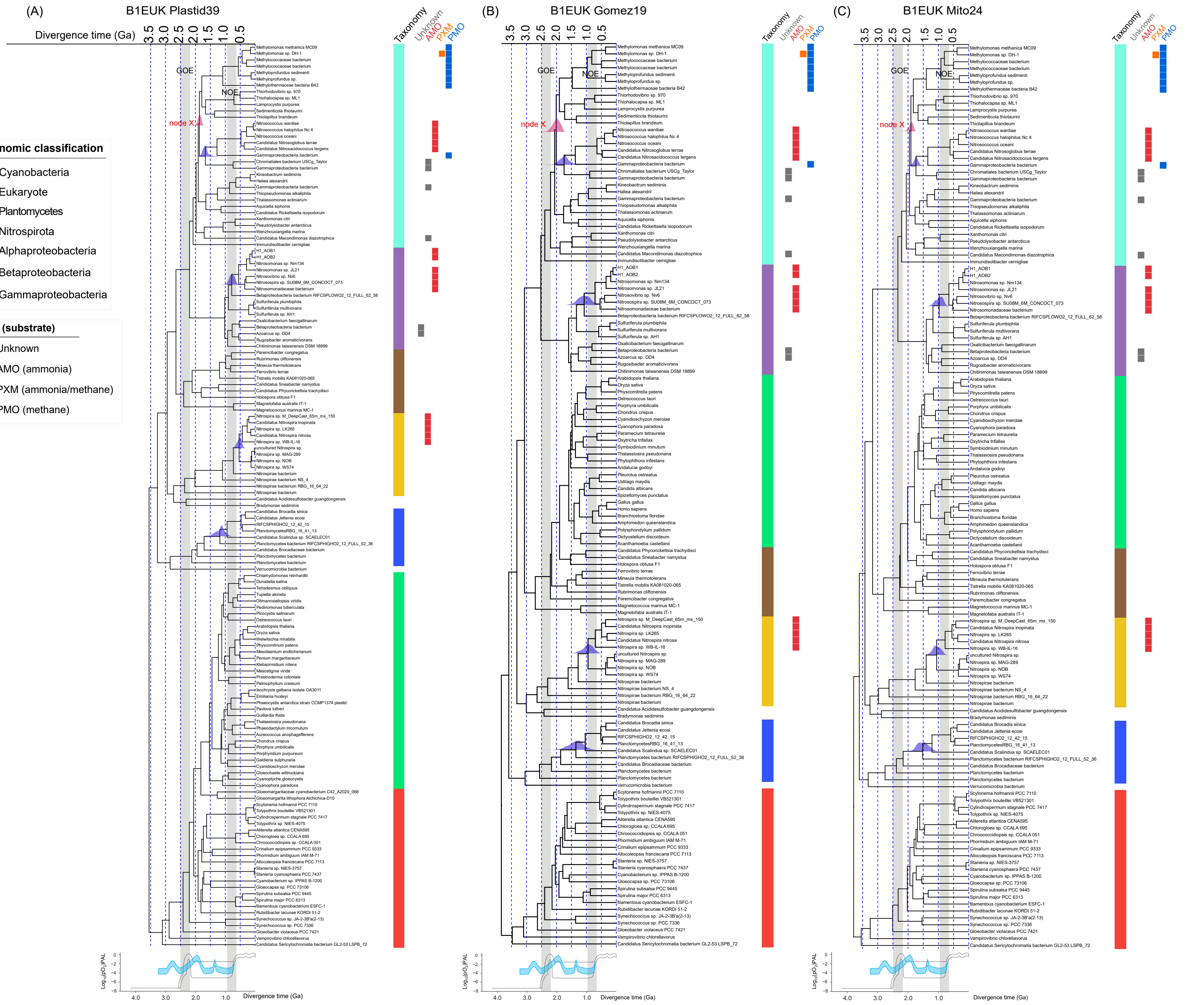


**Figure S3**. The chronogram of AOB and anammox bacteria estimated by sequential dating analysis. Each contains a timetree (upper) and the history of atmospheric oxygen levels of the Earth (lower). (A) The timetree was generated using the Plastid39 dataset, with calibrations (B1EUK) derived from the first-step of Bayesian sequential analysis and alignments of 15 plastid encoded proteins (9,598 sites). (B) The timetree was generated using the Gomez19 dataset, with calibrations (B1EUK) from the first-step of Bayesian sequential analysis and alignments of 13 eukaryotic nuclear proteins (13,118 sites). (C) The timetree was generated using the Mito24 dataset, with calibrations (B1EUK) from the first-step of Bayesian sequential analysis and alignments of six mitochondrial proteins (1,689 sites). The node X represents the LCA of Gamma-MOB and Gamma-AOB. The vertical grey bars correspond to the periods of the GOE and the NOE. The color strip adjacent to the label represents the major lineages of the analyzed genomes. The squares denote the XMO types found in the xmo-containing genomes used in molecular dating analysis. The density plot (shown in blue) on the nodes corresponding to the total groups of Gamma-AOB, of Beta-AOB, of CAOB, and of anammox bacteria, respectively, represent the posterior age estimates. The dark dashed line represents an oxygen model proposed by Lyons et al. (2014). The blue filled curve represents a more recent inference of atmospheric oxygen levels during the Archean and Proterozoic based on six geochemical proxies (Large et al. 2022). *$pO_2$*, atmospheric partial pressure of oxygen relative to the present atmospheric level (PAL).

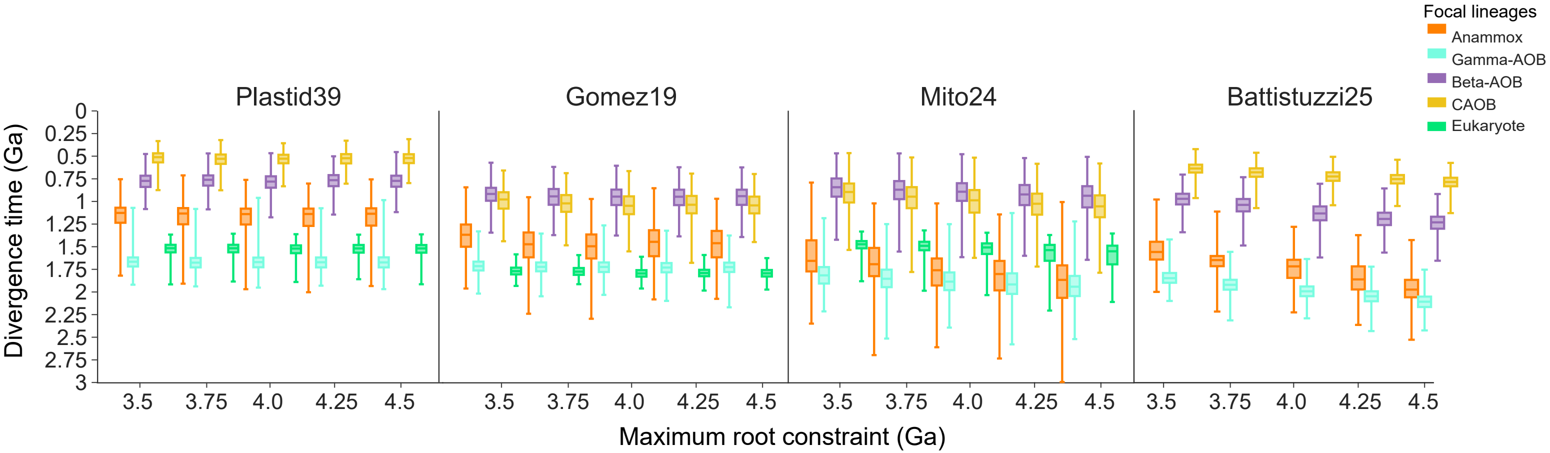


**Figure S4**. The posterior ages of five focal lineages based on the Bayesian sequential dating analysis using the same eukaryotic calibrations and bacterial calibrations (B1 and EUK in Dataset S2.1) and the different root maximum calibrations (from 3.5 to 4.5 Ga). For each gene set, we used the same eukaryotic calibrations derived from the first-step of the sequential dating analysis (see Supplementary Text Section 3.5; Figure 3). For the Battistuzzi25 gene set, it uses traditional dating analysis and only bacterial calibrations.

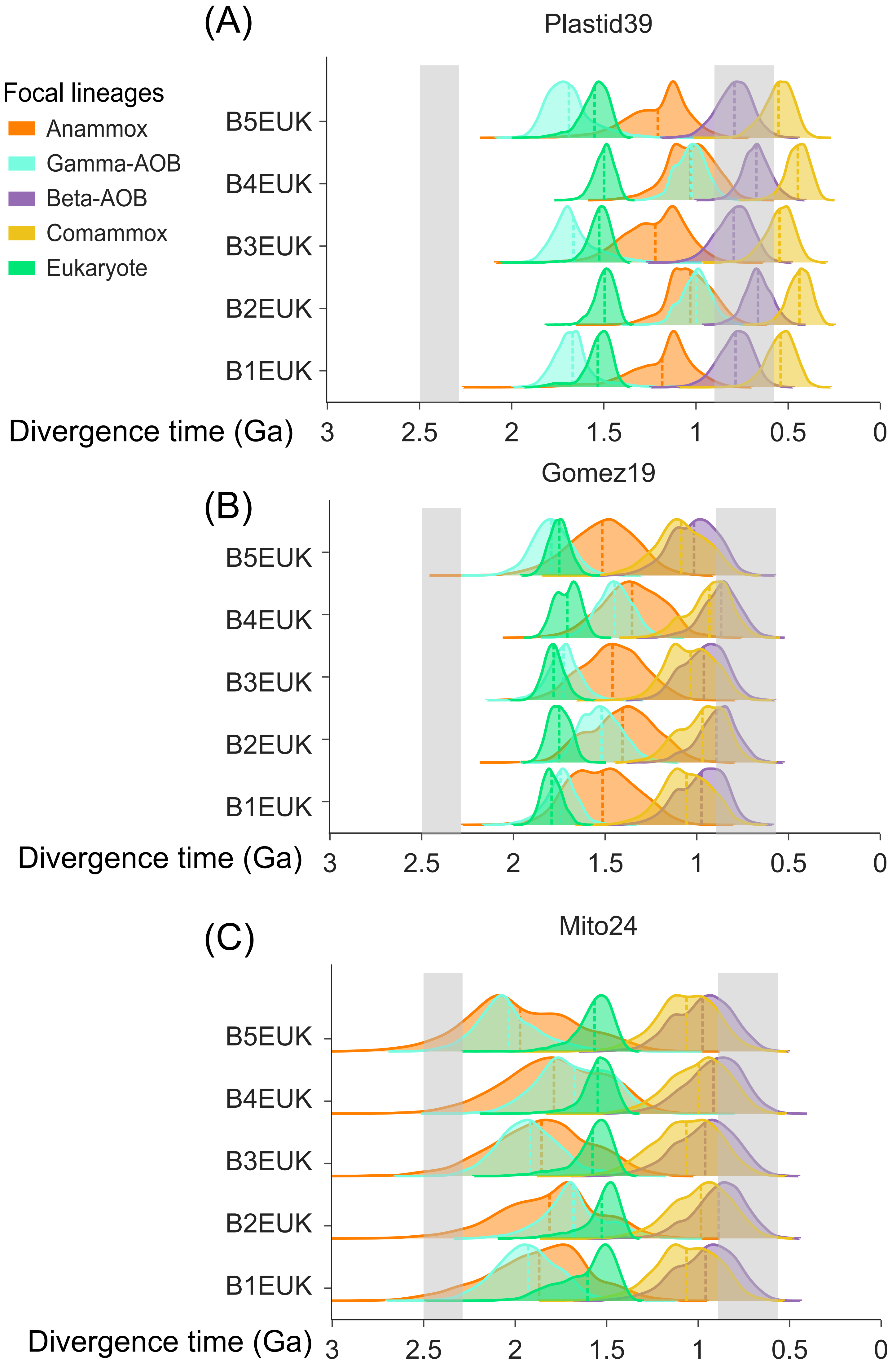


**Figure S5**. The posterior ages of five focal lineages which were estimated by the Bayesian sequential dating analysis using the same eukaryotic calibrations and different bacterial calibrations (B1EUK to B5EUK in Dataset S2.1). For each gene set, we implemented the same eukaryotic calibrations derived from the first-step of the sequential dating analysis (see Supplementary Text Section 3.5; Figure 3). The posterior time estimates are provided in the dataset S2.3.

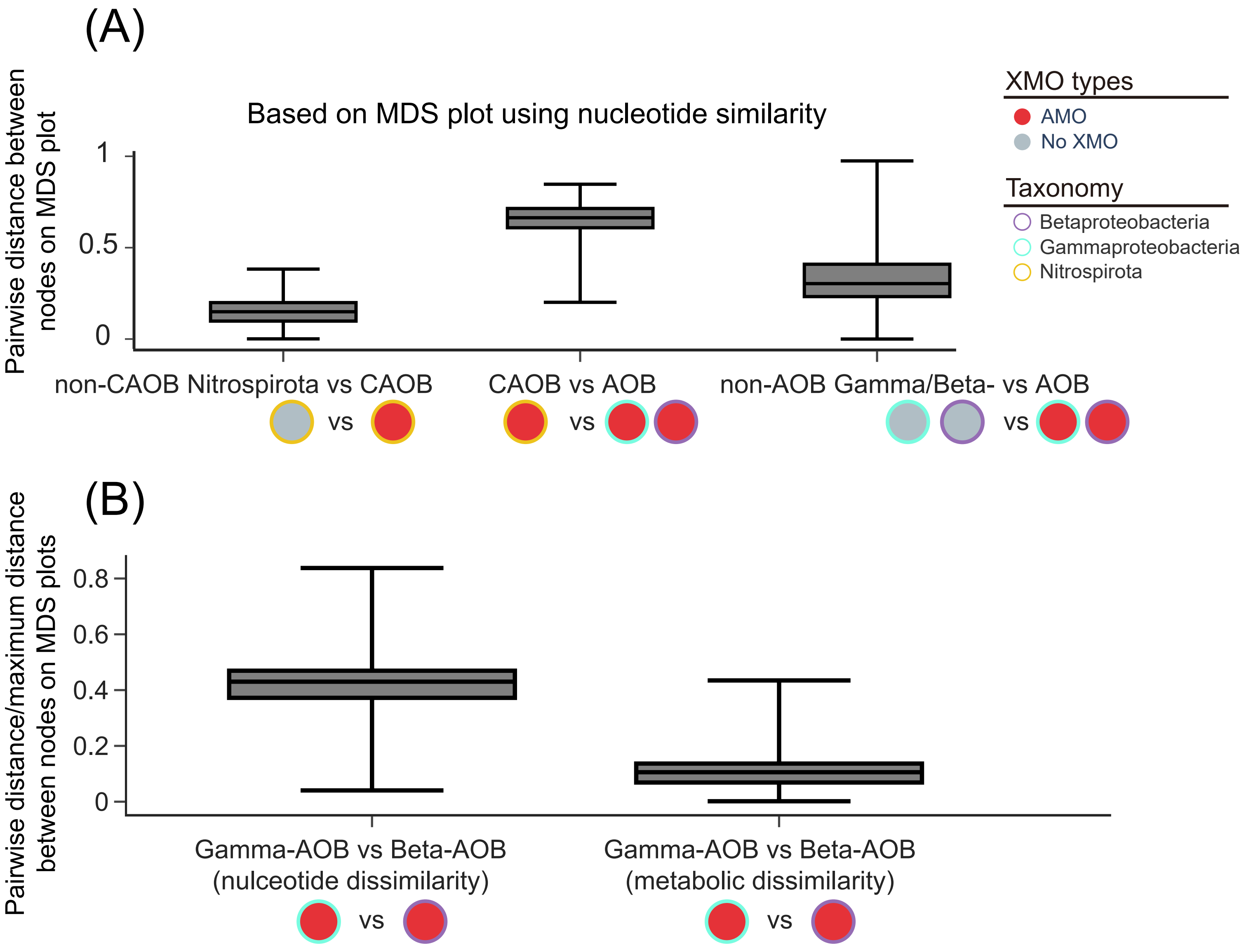


**Figure S6.** The pairwise distance between nodes on MDS plots. (A) The pairwise distance calculated based on the MDS plot, which indicates the nucleotide dissimilarities between genomes. The three box plots from left to right indicate the genetic distances between CAOB and non-CAOB Nitrospirota, between CAOB and AOB, and between AOB and non-AOB Gamma/Betaproteobacteria. (B) The normalized pairwise distance between Gamma-AOB and Beta-AOB was calculated based on two different MDS plots. The genetic distance was normalized by the maximum distance in the corresponding MDS plot. The left and the right box plot represent the nucleotide dissimilarities and the metabolic dissimilarities, respectively. The nodes shown below the x-axis indicate the compared groups shown in Figure 4.

### References


Álvarez-Carretero S, Tamuri AU, Battini M, Nascimento FF, Carlisle E, Asher RJ, Yang Z, Donoghue PC, Dos Reis M. 2022. A species-level timeline of mammal evolution integrating phylogenomic data. Nature 602:263-267.

Amard B, Bertrand-Sarfati J. 1997. Microfossils in 2000 Ma old cherty stromatolites of the Franceville Group, Gabon. Precambrian Research 81:197-221.

Angelis K, Alvarez-Carretero S, Dos Reis M, Yang Z. 2018. An Evaluation of Different Partitioning Strategies for Bayesian Estimation of Species Divergence Times. Syst Biol 67:61-77.

Aramaki T, Blanc-Mathieu R, Endo H, Ohkubo K, Kanehisa M, Goto S, Ogata H. 2020. KofamKOALA: KEGG Ortholog assignment based on profile HMM and adaptive score threshold. Bioinformatics 36:2251-2252.

Arp DJ, Chain PS, Klotz MG. 2007. The impact of genome analyses on our understanding of ammonia-oxidizing bacteria. Annu. Rev. Microbiol. 61:503-528.

Banerjee R, Proshlyakov Y, Lipscomb JD, Proshlyakov DA. 2015. Structure of the key species in the enzymatic oxidation of methane to methanol. Nature 518:431-434.

Barry P, Taylor L. 2013. Age of the Earth. In: Rink WJ, Thompson J, editors. Encyclopedia of Scientific Dating Methods. Dordrecht: Springer Netherlands. p. 1-2.

Battistuzzi FU, Hedges SB. 2009. A major clade of prokaryotes with ancient adaptations to life on land. Mol Biol Evol 26:335-343.

Bekker A, Holland HD, Wang PL, Rumble D, 3rd, Stein HJ, Hannah JL, Coetzee LL, Beukes NJ. 2004. Dating the rise of atmospheric oxygen. Nature 427:117-120.

Benson DA, Cavanaugh M, Clark K, Karsch-Mizrachi I, Lipman DJ, Ostell J, Sayers EW. 2013. GenBank. Nucleic Acids Res 41:D36-42.

Bergsten J. 2005. A review of long-branch attraction. Cladistics 21:163-193.

Blomberg SP, Garland Jr T, Ives AR. 2003. Testing for phylogenetic signal in comparative data: behavioral traits are more labile. Evolution 57:717-745.

Boddicker AM, Mosier AC. 2018. Genomic profiling of four cultivated Candidatus Nitrotoga spp. predicts broad metabolic potential and environmental distribution. ISME J 12:2864-2882.

Brocks JJ, Love GD, Summons RE, Knoll AH, Logan GA, Bowden SA. 2005. Biomarker evidence for green and purple sulphur bacteria in a stratified Palaeoproterozoic sea. Nature 437:866-870.

Coleman NV, Le NB, Ly MA, Ogawa HE, McCarl V, Wilson NL, Holmes AJ. 2012. Hydrocarbon monooxygenase in Mycobacterium: recombinant expression of a member of the ammonia monooxygenase superfamily. ISME J 6:171-182.

Cutting S, Roels S, Losick R. 1991. Sporulation operon spoIVF and the characterization of mutations that uncouple mother-cell from forespore gene expression in Bacillus subtilis. Journal of molecular biology 221:1237-1256.

Daims H, Lebedeva EV, Pjevac P, Han P, Herbold C, Albertsen M, Jehmlich N, Palatinszky M, Vierheilig J, Bulaev A, et al. 2015. Complete nitrification by Nitrospira bacteria. Nature 528:504-509.

Delignette-Muller ML, Dutang C. 2015. fitdistrplus: An R package for fitting distributions.

Journal of statistical software 64:1-34.

Dos Reis M, J I, M H, R. J. A, PCJ D, Z Y. 2012. Phylogenomic datasets provide both precision and accuracy in estimating the timescale of placental mammal phylogeny. Proc Biol Sci 279:3491-3500.

El-Gebali S, Mistry J, Bateman A, Eddy SR, Luciani A, Potter SC, Qureshi M, Richardson LJ, Salazar GA, Smart A, et al. 2019. The Pfam protein families database in 2019. Nucleic Acids Res 47:D427-D432.

Ester M, Kriegel H, Sander J, Xiaowei X. 1996. A density-based algorithm for discovering clusters in large spatial databases with noise. In: AAAI Press, Menlo Park, CA (United States).

Fu LM, Niu BF, Zhu ZW, Wu ST, Li WZ. 2012. CD-HIT: accelerated for clustering the next-generation sequencing data. Bioinformatics 28:3150-3152.

Golubic S, Lee SJ. 1999. Early cyanobacterial fossil record: preservation, palaeoenvironments and identification. European Journal of Phycology 34:339-348.

Golubic S, Sergeev VN, Knoll AH. 1995. Mesoproterozoic Archaeoellipsoides: Akinetes of heterocystous cyanobacteria. Lethaia 28:285-298.

Gonzalez-Cabaleiro R, Curtis TP, Ofiteru ID. 2019. Bioenergetics analysis of ammonia-oxidizing bacteria and the estimation of their maximum growth yield. Water Res 154:238-245.

Green AR, Hayes RP, Xun L, Kang C. 2012. Structural understanding of the glutathione-dependent reduction mechanism of glutathionyl-hydroquinone reductases. J Biol Chem 287:35838-35848.

Harden MM, He A, Creamer K, Clark MW, Hamdallah I, Martinez KA, 2nd, Kresslein RL, Bush SP, Slonczewski JL. 2015. Acid-adapted strains of Escherichia coli K-12 obtained by experimental evolution. Appl Environ Microbiol 81:1932-1941.

Horodyski RJ, Donaldson JA. 1980. Microfossils from the middle Proterozoic Dismal Lakes groups, arctic Canada. Precambrian Research 11:125-159.

Hugoson E, Lam WT, Guy L. 2020. miComplete: weighted quality evaluation of assembled microbial genomes. Bioinformatics 36:936-937.

Hutter B, Dick T. 1998. Increased alanine dehydrogenase activity during dormancy in Mycobacterium smegmatis. FEMS Microbiology Letters 167:7-11.

Johnson M, Zaretskaya I, Raytselis Y, Merezhuk Y, McGinnis S, Madden TL. 2008. NCBI BLAST: a better web interface. Nucleic Acids Res 36:W5-9.

Kalyaanamoorthy S, Minh BQ, Wong TKF, von Haeseler A, Jermiin LS. 2017. ModelFinder: fast model selection for accurate phylogenetic estimates. Nat Methods 14:587-589.

Kanehisa M, Goto S. 2000. KEGG: Kyoto Encyclopedia of Genes and Genomes. Nucleic Acids Research 28:27-30.

Kans J. 2020. Entrez direct: E-utilities on the UNIX command line. In. Entrez Programming Utilities Help [Internet]: National Center for Biotechnology Information (US).

Katoh K, Standley DM. 2013. MAFFT multiple sequence alignment software version 7: improvements in performance and usability. Mol Biol Evol 30:772-780.

Khadka R, Clothier L, Wang L, Lim CK, Klotz MG, Dunfield PF. 2018. Evolutionary History of Copper Membrane Monooxygenases. Front Microbiol 9:2493.

Kishino H, Miyata T, Hasegawa M. 1990. Maximum likelihood inference of protein phylogeny

and the origin of chloroplasts. Journal of Molecular Evolution 31:151-160.
Klotz MG, Schmid MC, Strous M, op den Camp HJ, Jetten MS, Hooper AB. 2008. Evolution of an octahaem cytochrome c protein family that is key to aerobic and anaerobic ammonia oxidation by bacteria. Environ Microbiol 10:3150-3163.
Klotz MG, Stein LY. 2008. Nitrifier genomics and evolution of the nitrogen cycle. FEMS Microbiol Lett 278:146-156.
Kozlowski JA, Kits KD, Stein LY. 2016. Comparison of nitrogen oxide metabolism among diverse ammonia-oxidizing bacteria. Front Microbiol 7:1090.
Lehtovirta-Morley LE. 2018. Ammonia oxidation: Ecology, physiology, biochemistry and why they must all come together. Fems Microbiology Letters 365.
Letunic I, Bork P. 2021. Interactive Tree Of Life (iTOL) v5: an online tool for phylogenetic tree display and annotation. Nucleic Acids Res 49:W293-W296.
Liao T, Wang S, Stueken EE, Luo H. 2022. Phylogenomic Evidence for the Origin of Obligate Anaerobic Anammox Bacteria Around the Great Oxidation Event. Mol Biol Evol 39.
Mai U, Sayyari E, Mirarab S. 2017. Minimum variance rooting of phylogenetic trees and implications for species tree reconstruction. PloS one 12:e0182238.
Matheus Carnevali PB, Schulz F, Castelle CJ, Kantor RS, Shih PM, Sharon I, Santini JM, Olm MR, Amano Y, Thomas BC, et al. 2019. Author Correction: Hydrogen-based metabolism as an ancestral trait in lineages sibling to the Cyanobacteria. Nat Commun 10:1451.
Minh BQ, Nguyen MA, von Haeseler A. 2013. Ultrafast approximation for phylogenetic bootstrap. Mol Biol Evol 30:1188-1195.
Mistry J, Finn RD, Eddy SR, Bateman A, Punta M. 2013. Challenges in homology search: HMMER3 and convergent evolution of coiled-coil regions. Nucleic Acids Res 41:e121.
Munoz-Gomez SA, Hess S, Burger G, Lang BF, Susko E, Slamovits CH, Roger AJ. 2019. An updated phylogeny of the Alphaproteobacteria reveals that the parasitic Rickettsiales a Holosporales have independent origins. Elife 8.
Munoz-Gomez SA, Susko E, Williamson K, Eme L, Slamovits CH, Moreira D, Lopez-Garcia P, Roger AJ. 2022. Site-and-branch-heterogeneous analyses of an expanded dataset favour mitochondria as sister to known Alphaproteobacteria. Nat Ecol Evol 6:253-262.
Nguyen LT, Schmidt HA, von Haeseler A, Minh BQ. 2015. IQ-TREE: a fast and effective stochastic algorithm for estimating maximum-likelihood phylogenies. Mol Biol Evol 32:268-274.
Ondov BD, Treangen TJ, Melsted P, Mallonee AB, Bergman NH, Koren S, Phillippy AM. 2016. Mash: fast genome and metagenome distance estimation using MinHash. Genome Biol 17:132.
Parks DH, Rinke C, Chuvochina M, Chaumeil PA, Woodcroft BJ, Evans PN, Hugenholtz P, Tyson GW. 2017. Recovery of nearly 8,000 metagenome-assembled genomes substantially expands the tree of life. Nat Microbiol 2:1533-1542.
Philippe H, Brinkmann H, Lavrov DV, Littlewood DTJ, Manuel M, Wörheide G, Baurain D. 2011. Resolving difficult phylogenetic questions: why more sequences are not enough. PLoS biology 9:e1000602.
Picone N, Pol A, Mesman R, van Kessel M, Cremers G, van Gelder AH, van Alen TA, Jetten MSM, Lucker S, Op den Camp HJM. 2021. Ammonia oxidation at pH 2.5 by a new

gammaproteobacterial ammonia-oxidizing bacterium. ISME J 15:1150-1164.

Ponce-Toledo RI, Deschamps P, López-García P, Zivanovic Y, Benzerara K, Moreira D. 2017. An early-branching freshwater cyanobacterium at the origin of plastids. Current Biology 27:386-391.

Quang le S, Gascuel O, Lartillot N. 2008. Empirical profile mixture models for phylogenetic reconstruction. Bioinformatics 24:2317-2323.

Rannala B, Yang Z. 2007. Inferring speciation times under an episodic molecular clock. Syst Biol 56:453-466.

Revell LJ. 2012. phytools: an R package for phylogenetic comparative biology (and other things). Methods in ecology and evolution:217-223.

Sander C, Schneider R. 1991. Database of homology - derived protein structures and the structural meaning of sequence alignment. Proteins: Structure, Function, and Bioinformatics 9:56-68.

Sanderson MJ, Baldwin B, Bharathan G, Campbell C, Von Dohlen C, Ferguson D, Porter J, Wojciechowski M, Donoghue M. 1993. The growth of phylogenetic information and the need for a phylogenetic data base. Syst Biol 42:562-568.

Sarkar S, Bhattacharyya A, Mandal S. 2015. YnfA, a SMR family efflux pump is abundant in Escherichia coli isolates from urinary infection. Indian journal of medical microbiology 33:139-142.

Schubert E, Sander J, Ester M, Kriegel HP, Xu X. 2017. DBSCAN revisited, revisited: why and how you should (still) use DBSCAN. ACM Transactions on Database Systems (TODS) 42:1-21.

Seemann T. 2014. Prokka: rapid prokaryotic genome annotation. Bioinformatics 30:2068-2069.

Segata N, Bornigen D, Morgan XC, Huttenhower C. 2013. PhyloPhlAn is a new method for improved phylogenetic and taxonomic placement of microbes. Nat Commun 4:2304.

Shen X-X, Hittinger CT, Rokas A. 2017. Contentious relationships in phylogenomic studies can be driven by a handful of genes. Nat Ecol Evol 1:0126.

Shimodaira H. 2002. An approximately unbiased test of phylogenetic tree selection. Syst Biol 51:492-508.

Smith SA, Walker-Hale N, Walker JF, Brown JW. 2020. Phylogenetic Conflicts, Combinability, and Deep Phylogenomics in Plants. Syst Biol 69:579-592.

Steenwyk JL, Buida TJ, Li YN, Shen XX, Rokas A. 2020. ClipKIT: A multiple sequence alignment trimming software for accurate phylogenomic inference. Plos Biology 18.

Stein LY. 2019. Insights into the physiology of ammonia-oxidizing microorganisms. Curr Opin Chem Biol 49:9-15.

Strassert JFH, Irisarri I, Williams TA, Burki F. 2021. Author Correction: A molecular timescale for eukaryote evolution with implications for the origin of red algal-derived plastids. Nat Commun 12:3574.

Tavormina PL, Orphan VJ, Kalyuzhnaya MG, Jetten MS, Klotz MG. 2011. A novel family of functional operons encoding methane/ammonia monooxygenase-related proteins in gammaproteobacterial methanotrophs. Environ Microbiol Rep 3:91-100.

Van Kranendonk MJ. 2006. Volcanic degassing, hydrothermal circulation and the flourishing of early life on Earth: A review of the evidence from c. 3490-3240 Ma rocks of the Pilbara Supergroup, Pilbara Craton, Western Australia. Earth-Science Reviews 74:197-240.

Wang HC, Minh BQ, Susko E, Roger AJ. 2018. Modeling Site Heterogeneity with Posterior Mean Site Frequency Profiles Accelerates Accurate Phylogenomic Estimation. Syst Biol 67:216-235.

Wang S, Luo H. 2023. Dating the bacterial tree of life based on ancient symbiosis. bioRxiv:2023.2006.2018.545440.

Wang SS, Luo HW. 2021. Dating Alphaproteobacteria evolution with eukaryotic fossils. Nature Communications 12.

Wang Z, Wu M. 2015. An integrated phylogenomic approach toward pinpointing the origin of mitochondria. Scientific reports 5:7949.

Yang Z, Rannala B. 2006. Bayesian estimation of species divergence times under a molecular clock using multiple fossil calibrations with soft bounds. Mol Biol Evol 23:212-226.

Yang ZH. 2007. PAML 4: Phylogenetic analysis by maximum likelihood. Molecular Biology and Evolution 24:1586-1591.

Yin QJ, Zhang WJ, Qi XQ, Zhang SD, Jiang T, Li XG, Chen Y, Santini CL, Zhou H, Chou IM, et al. 2017. High Hydrostatic Pressure Inducible Trimethylamine N-Oxide Reductase Improves the Pressure Tolerance of Piezosensitive Bacteria Vibrio fluvialis. Front Microbiol 8:2646.

Zhang H, Sun Y, Zeng Q, Crowe SA, Luo H. 2021. Snowball Earth, population bottleneck and Prochlorococcus evolution. Proc Biol Sci 288:20211956.

Zhang H, Wang S, Liao T, Crowe SA, Luo H. 2023. Emergence of Prochlorococcus in the Tonian oceans and the initiation of Neoproterozoic oxygenation. bioRxiv:2023.2009.2006.556545.

Zhu Q, Mai U, Pfeiffer W, Janssen S, Asnicar F, Sanders JG, Belda-Ferre P, Al-Ghalith GA, Kopylova E, McDonald D, et al. 2019. Phylogenomics of 10,575 genomes reveals evolutionary proximity between domains Bacteria and Archaea. Nat Commun 10:5477.
